# Epigenetic maintenance of PRC2-repressed chromatin requires RTT109 but not H3K56 acetylation

**DOI:** 10.64898/2026.05.14.724656

**Authors:** Rochelle E. Yap, Felicia Ebot-Ojong, Abigail J. Ameri-Solanky, Zachary A. Lewis

**Author notes:** Corresponding Author: Zachary A. Lewis, University of Georgia, Athens, GA.

## Abstract

In animals, plants, and some fungi, Polycomb Repressive Complex 2 (PRC2) catalyzes trimethylation of histone H3 lysine 27 (H3K27me3) to establish transcriptionally repressed chromatin. Here, we identify the histone acetyltransferase RTT109 as a key regulator of PRC2-repressed domains in the model fungus *Neurospora crassa*. Although RTT109 interacts with the VPS75 homolog Nucleosome Assembly Factor 2 (NAF-2), we show that proper structure and function of PRC2-methylated chromatin require RTT109 catalytic activity but are independent of NAF-2 and H3K56 acetylation. We further demonstrate that H3K27me3 can be stably propagated over multiple rounds of mitosis in the absence of sequence-specific PRC2 targeting, and that RTT109 is essential for maintenance of the repressed state. These findings uncover a replication-linked mechanism for epigenetic memory and establish RTT109 as a key regulator of Polycomb-mediated chromatin inheritance.

## INTRODUCTION

In eukaryotes, histone post-translational modifications (PTMs) directly influence transcriptional regulation by altering DNA accessibility and controlling the landscape of chromatin interacting proteins^1^. Some histone PTMs are stably propagated over mitosis and contribute to epigenetic maintenance of gene expression programs^2^. For example, Polycomb Repressive Complex 1 (PRC1) and PRC2 catalyze mono-ubiquitylation on Histone H2A and methylation of Histone H3 Lysine 27 (H3K27me), respectively, to assemble stably repressed chromatin domains, often referred to as facultative heterochromatin^3,4^. In mammals and plants, the PRCs are essential for maintaining cell identity by repressing genes that specify alternative cell fates^5,6^. In some fungi, PRC2 components are present and have been linked to multicellular development, stress responses, and virulence^7–10^. While many fundamental aspects of Polycomb-mediated repression are conserved, the mechanisms that establish and maintain facultative heterochromatin vary between species, highlighting major gaps in our understanding of chromatin regulation across eukaryotes.

In metazoans, cell-type specific patterns of H3K27me3 patterns are established during development. In *Drosophila melanogaster*, where PRCs were originally discovered, cis-regulatory DNA elements known as Polycomb Response Elements recruit PRC1, PRC2, or accessory proteins to direct deposition of H3K27me3^5,6,11^. PRCs are recruited in a similar manner to CpG islands in mammals and to discrete DNA binding sites in plants^12,13^. Once established, H3K27me3 is propagated along the chromatin fiber via the read-write activity of PRC2^14,15^. The PRC2 subunit embryonic ectoderm development (EED) binds to H3K27me3 (“read”) and allosterically stimulates PRC2 methyltransferase activity (“write”), often leading to the formation of large, repressed domains that encompass multiple genes. Notably, genes within these domains can remain stably methylated across multiple rounds of mitosis in the absence of continuous PRC recruitment^16,17^. This long-term stability is remarkable given that modified histones are passively diluted each replication cycle^2^. In plants and animals, mitotic inheritance of the PRC-repressed chromatin state depends on a variety of mechanisms including positive feedback between the PRCs and replication-associated proteins^18–24^, but the genes and mechanisms that underlie this epigenetic memory are still being elucidated.

Many filamentous fungi assemble PRC2-dependent facultative heterochromatin, but PRC1 was lost in an early fungal ancestor^25^. In the model filamentous fungus *Neurospora crassa*, H3K27me3 is enriched across large, multi-gene domains that are clustered together within the three-dimensional space of the nucleus^26–29^. PRC2-methylated genes are expressed during multicellular fruiting body development but are stably repressed in vegetative tissues (mycelium)^30^. Thus, *N. crassa* is an attractive experimental system to investigate the control and function of PRC2. Three core PRC2 components, SET-7 (homolog of Enhancer of zeste; HsEZH1/2), EED, and SU(Z)12 (homolog of suppressor of zeste-12), are required for H3K27me3, as in other eukaryotes^26^. H3K27me3 in *N. crassa* is directed by telomere repeats (TTAGGGn) and by the PRC2 Associated Subunit (PAS) protein^31,32^. Other regulators required for steady-state H3K27me3 levels and gene repression include the chromatin remodeler Imitation Switch (ISW), NPF/CAC-3, hH2A.Z, and components of constitutive heterochromatin^26,33–37^. Finally, members of the RPD3L (Reduced Potassium Dependency 3 Large) histone deacetylase complex and EPR-1 (Effector of Polycomb Repression 1) are required for Polycomb-mediated repression without concurrent loss of H3K27me3^28,38^.

In several filamentous fungi including *N. crassa,* H3K27me3 regions are co-enriched with H3K36me3 deposited by ASH1 (homolog of Absent, Small, or Homeotic Disc 1)^39,40^. Notably, the H3K36me3 enrichment pattern in facultative heterochromatin is distinct from that of euchromatin. In facultative heterochromatin, H3K36me3 is enriched across large, multi-gene domains covering both promoters and gene bodies^39^. In euchromatin, however, H3K36me3 is deposited co-transcriptionally by SET-2 and therefore restricted to coding sequences of expressed genes^39^. Loss of ASH1-catalyzed H3K36me3 leads to defective gene silencing and altered H3K27me3 patterns^39,41^. This differs from the situation in animals, where Ash1 and H3K36me3 are reported to antagonize H3K27me3 *in vivo* and *in vitro*^13^. Although genetic studies have uncovered genes that are required for normal structure and function of facultative heterochromatin in *N. crassa,* much remains unknown. For example, can the PRC2-repressed chromatin state be stably transmitted across mitosis in the absence of a PRC1 homolog, and if so, do replication associated-factors contribute to epigenetic stability?

Here, we show the histone acetyltransferase <u>R</u>egulator of <u>T</u>y1 transposition 109 (RTT109) is required for stable facultative heterochromatin in *N. crassa*. RTT109 is a fungal-specific Histone H3 Lysine 56 (H3K56) acetyltransferase originally discovered in *Saccharomyces cerevisiae* and has previously been implicated in DNA replication, DNA-repair and transcriptional control^42^. RTT109 is responsible for H3K56 acetylation (H3K56ac) on newly synthesized H3 during replication-coupled nucleosome assembly (RCNA), prior to histone deposition by Chromatin Assembly Factor Complex (CAF-1)^43^. In yeast, two histone chaperones, Vacuolar Protein Sorting (Vps75) and Anti-silencing Factor (Asf1) are important for RTT109’s catalytic activity^44^. In addition to its established role in replication, *N. crassa* RTT109 is linked to small RNA production, quelling, and homologous recombination^45^. Here, we report that RTT109 regulates facultative heterochromatin in *N. crassa* independent of H3K56 acetylation. We demonstrate that H3K27me3 is stably maintained in the absence of continuous PRC2 recruitment, and we show that RTT109 is required for stable mitotic inheritance of H3K27me3. These data provide new insights into the genes and mechanisms responsible for epigenome stability in eukaryotes.

## RESULTS

### RTT109 is required for repression of a subset of facultative heterochromatin genes

We screened an RNA-seq dataset to find gene knockout strains with defective repression of H3K27me3-marked genes, identifying RTT109 as a candidate regulator of facultative heterochromatin in *N. crassa*^46^. To confirm the *rtt109* phenotype, we deleted *rtt109* in a reporter strain harboring a silent basta resistance gene (*bar*) embedded within a facultative heterochromatin domain (*ncu06889::bar^OFF^*). Deletion of *rtt109* produced a basta-resistant phenotype, similar to the *Δeed* control strain lacking a functional PRC2 complex (Fig. 1a). We next analyzed RNA-seq data to compare global gene expression in mycelia of wild type, Δ*rtt109,* and a Δ*set-7* control strain. As expected, PRC2-methylated genes (n=573) were tightly repressed in wild type. In contrast, the average transcript levels of this gene set were significantly increased for both Δ*rtt109* and the Δ*set-7* control strain (Supplementary Fig. 1a, Supplementary Data Tab 1). To determine if the observed defects in facultative heterochromatin-associated gene repression are specific to Δ*rtt109* or a general response to hypoacetylation, we performed RNA-seq on mutants lacking other histone acetyltransferases. Broad upregulation of facultative heterochromatin genes was only observed in Δ*rtt109* (Supplementary Fig. 1b; Supplementary Data Tab 2). In total, 136 of 573 H3K27me3-enriched genes showed increased expression in Δ*rtt109* compared to wild type (Supplementary Fig. 1c). More PRC2-methylated genes were induced in *Δrtt109* than in the *Δset-7* control strain. Across all *N. crassa* genes, we found that 722 genes exhibited differential expression in *Δrtt109* compared to wild type, with 705/722 genes exhibiting increased expression (Fig. 1b, Supplementary Data Tab 3). Importantly, known components of the facultative heterochromatin pathway were expressed at normal levels in the Δ*rtt109* strain (Supplementary Data Tab 3).

**Figure 1.**
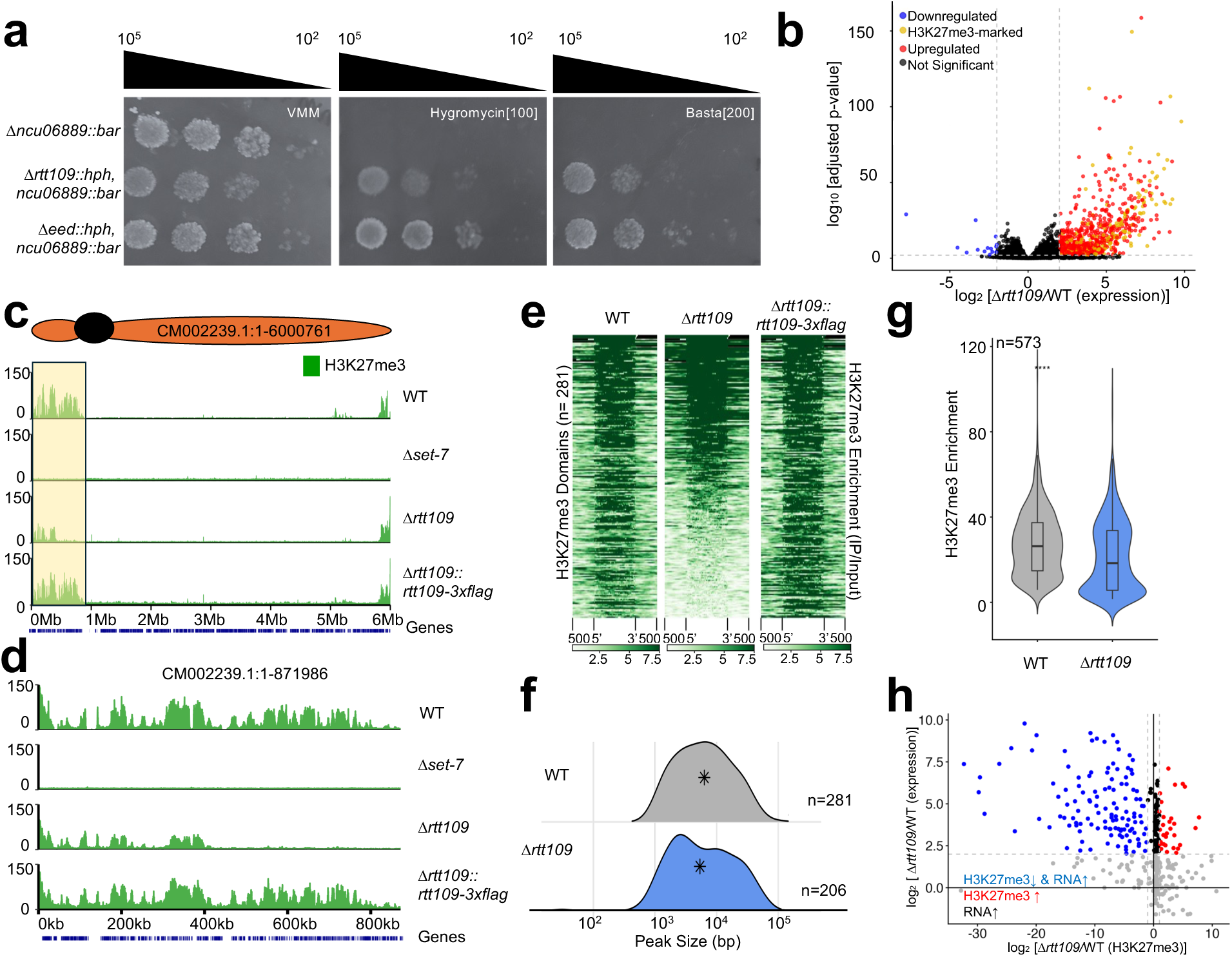
The *Δrtt109* strain exhibits defects in facultative heterochromatin. **a.** Deletion of *rtt109* leads to activation of a facultative heterochromatin reporter gene. Serial dilutions of conidia for the indicated strains were spotted on VMM, VMM+ hygromycin, and VMM+ BASTA. Growth on BASTA-containing medium indicates derepression of the normally silenced Δ*ncu06889::bar^OFF^* reporter. The number of cells per spot is indicated above each panel. **b.** The volcano plot shows relative expression levels (x-axis; log_2_ [*Δrtt109/*WT]) and log transformed adjusted p-values (y-axis) for all *N. crassa* genes. Genes that passed fold-change and statistical thresholds (adjusted p-value < 0.01 and |log_2_ [*Δrtt109/*WT]| > 2) are colored by their expression pattern: down in Δ*rtt109* (blue), up in Δ*rtt109* (red), and upregulated H3K27me3-marked genes (yellow). Genes below the statistical threshold are colored in black. **c.** The genome browser tracks show relative enrichment of H3K27me3 (ChIP-seq) across LG IV for the indicated strains. Normalized coverage values are shown on the y-axis and chromosomal position is shown on the x-axis. The bottom track (blue) indicates positions of annotated genes. **d.** Relative enrichment of H3K27me3 is shown for the region highlighted in yellow in panel c. **e.** The heatmaps show enrichment of H3K27me3 for the indicated strains. Each row shows enrichment across an individual H3K27me3 domain identified in wild type ± 500bp (n=281). The size of each domain is scaled. The domains are sorted based on enrichment level in Δ*rtt109* from highest to lowest. **f.** The ridge plots show distribution of H3K27me3 peak sizes from WT (n=281) and *Δrtt109* (n=206). The number of peaks (density; y-axis) is plotted versus peak size (x-axis) for all H3K27me3 peaks called using MACS3. The black asterisks indicate the median peak size for each group. Peak sizes are plotted on a log_10_ scale. **g.** The violin plots show H3K27me3 signal across H3K27me3-marked genes (n=573) from 3 input-normalized WT and *Δrtt109* biological replicates. Within each violin, the boxplots indicate the middle 50% values, the center bolded line indicates the median, the box above is the 75^th^ percentile, and the box below is the 25^th^ percentile. Asterisks indicate significance level calculated by paired Wilcoxon signed rank tests (formatted p-value < 2e-16). **h.** The dot plot shows relative gene expression log_2_ [*Δrtt109/*WT] compared to relative H3K27me3-enrichment log_2_ [*Δrtt109/*WT] for H3K27me3-marked genes. Genes are colored based on combined changes in H3K27me3 and RNA expression in *Δrtt109*. The color of the dot represents the phenotype observed for an individual H3K27me3-marked gene (red: log_2_ [*Δrtt109/*WT(H3K27me3)] > 1 and log_2_ [*Δrtt109/*WT(expression)] > 2; blue: log_2_ [*Δrtt109/*WT(H3K27me3)] < –1 and log_2_ [*Δrtt109/*WT(expression)] > 2; black: log_2_ [*Δrtt109/*WT(H3K27me3)] < 1 or <-1 and log_2_ [*Δrtt109/*WT(expression)] >). Grey dots do not match any of the aforementioned categories.

Genes that were differentially expressed in Δ*rtt109* showed significant overlap with PRC2-repressed genes. Of the 722 genes that were differentially expressed in Δ*rtt109,* 178 were also significantly upregulated in Δ*set-7,* and 78 of these genes were H3K27me3-marked (Supplementary Fig. 1c, d). Furthermore, the Δ*rtt109* strain mis-expressed genes and gene categories that are typically induced during fruiting body development, as previously reported for PRC2-deficient mutants^30^. StringGo enrichment analysis of the 705 genes upregulated in *Δrtt109* identified functional categories enriched during fruiting body development, including fungal cell wall synthesis and organization (Supplementary Fig. 1e, Supplementary Data Tab 4)^30^. Genes in the cell wall chitin metabolism pathway had the greatest proportion of significantly upregulated genes (71%; 15/21). We cross-referenced the list of upregulated genes in *Δrtt109* with a list of developmentally induced genes with and without H3K27me3. 600/722 genes upregulated in *Δrtt109* are also developmentally induced (Supplementary Fig. 1f, Supplementary Data Tab 5). This is reminiscent of the Δ*set-7* mutant, which forms aberrant fruiting bodies and exhibits broad upregulation of developmentally induced genes^30^. Together, these data show that RTT109 is required for normal gene repression in *N. crassa*, particularly within facultative heterochromatin domains.

### RTT109 is required for normal facultative heterochromatin formation

To determine whether transcriptional derepression in *Δrtt109* is accompanied by altered H3K27me3, we performed ChIP-seq to examine patterns of H3K27me3 across the *N. crassa* genome. Inspection of H3K27me3-enrichment patterns in the IGV genome browser revealed regional losses of H3K27me3 in the *Δrtt109* mutant. For example, we observed a complete loss of H3K27me3 in a ∼372kb region containing 39 H3K27me3-marked genes on Linkage Group IV (Fig. 1c, d). To confirm that altered H3K27me3 patterns were indeed caused by loss of RTT109, we performed complementation by integrating a *rtt109-3xflag* construct driven by the native promoter. Introduction of *rtt109-3xflag* restored H3K27me3 patterns (Fig. 1c, d). Loss of RTT109 has previously been shown to increase sensitivity to DNA-damaging agents such as methyl-methanesulfonate (MMS), reflective of its known role in chromatin assembly and DNA repair. Complementation also rescued the DNA-damage repair defect of Δ*rtt109* (Supplementary Fig. 2a). ChIP-seq results were confirmed by performing ChIP-qPCR (Supplementary Fig. 2b).

To quantify losses of H3K27me3, we used MACS3 software to identify H3K27me3-enriched regions in wild type and the Δ*rtt109* strain^47^. In wild type, 281 H3K27me3-enriched peaks corresponding to 7.13% of the *N. crassa* genome were identified, consistent with prior published data^36,48^. In the Δ*rtt109* strain, 206 H3K27me3-enriched regions covered ∼5.61% of the genome. 102 of 281 domains (33.6%) showed ≥50% loss of H3K27me3 signal in *Δrtt109* compared to wild type (Fig. 1e, Supplementary Data Tab 6). In addition to the reduced number of H3K27me3 domains, the enriched regions were smaller in *Δrtt109*, with the median peak size decreasing from 6.1 kb in wild type to 4.5 kb in *Δrtt109* (Fig. 1f). Overall, we observed strong regional losses of H3K27me3 in *Δrtt109* compared to wild type. At the gene level, the average H3K27me3 enrichment across 573 genes typically methylated by PRC2 was significantly reduced by 31% in *Δrtt109* (Fig. 1g; paired Wilcoxon signed-rank test, p < 0.001). In total, 204/573 H3K27me3-marked genes exhibited a ≥50% reduction in H3K27me3 enrichment level in *Δrtt109*. This loss is rescued in the *rtt109-3xflag* strain (Supplementary Fig. 2c, d).

Next, we asked how changes in H3K27me3 relate to transcriptional output at the individual gene level by plotting changes in expression against changes in H3K27me3 (Fig. 1h). We found that increased transcription was correlated with loss of H3K27me3, though 36/136 upregulated genes in Δ*rtt109* retained high H3K27me3 levels, suggesting the loss of H3K27me3 is not strictly required for transcriptional derepression in Δ*rtt109*.

We next asked if RTT109 is required for normal patterns of H3K36me3 in facultative heterochromatin. We performed ChIP-seq to examine the distribution of H3K36me3 across facultative heterochromatin domains for wild type, *Δrtt109,* Δ*set-7*, and the complemented *rtt109-3xflag* strain. The Δ*rtt109* strain displayed a marked reduction in H3K36me3 levels within facultative heterochromatin, but SET-2-dependent H3K36me3 in euchromatin was unaffected (Fig. 2a-b). In Δ*rtt109,* we observed 33% of H3K27me3-marked genes had significantly reduced H3K36me3 enrichment (Fig. 2d). These defects were rescued in the complemented strain (Fig. 2a, c-d). As previously reported, H3K36me3 patterns in facultative heterochromatin appeared largely unaffected in the Δ*set-7* strain (Fig. 2c-d)^39^. We next performed differential enrichment analysis to compare changes in H3K27me3 and H3K36me3 within facultative heterochromatin genes using a cut-off of |log_2_ fold change| ≥ 1. Across all facultative heterochromatin genes, the median H3K36me3 enrichment level was reduced from 6.4 to 4.19 in the Δ*rtt109* strain (Fig. 2e). Only a fraction of these genes exhibit loss of both H3K27me3 and H3K36me3 (40/573) using a fold change cutoff of –1. An additional 30/573 displayed reduced H3K36me3 without a corresponding loss of H3K27me3. Strikingly, 164/573 genes lose H3K27me3 without a corresponding loss of H3K36me3 (Fig. 2e). Together, these data suggest that H3K36me3 loss is not simply a secondary consequence of H3K27me3 loss, and reciprocally loss of H3K36me3 cannot explain the loss of H3K27me3 in *Δrtt109*.

**Figure 2.**
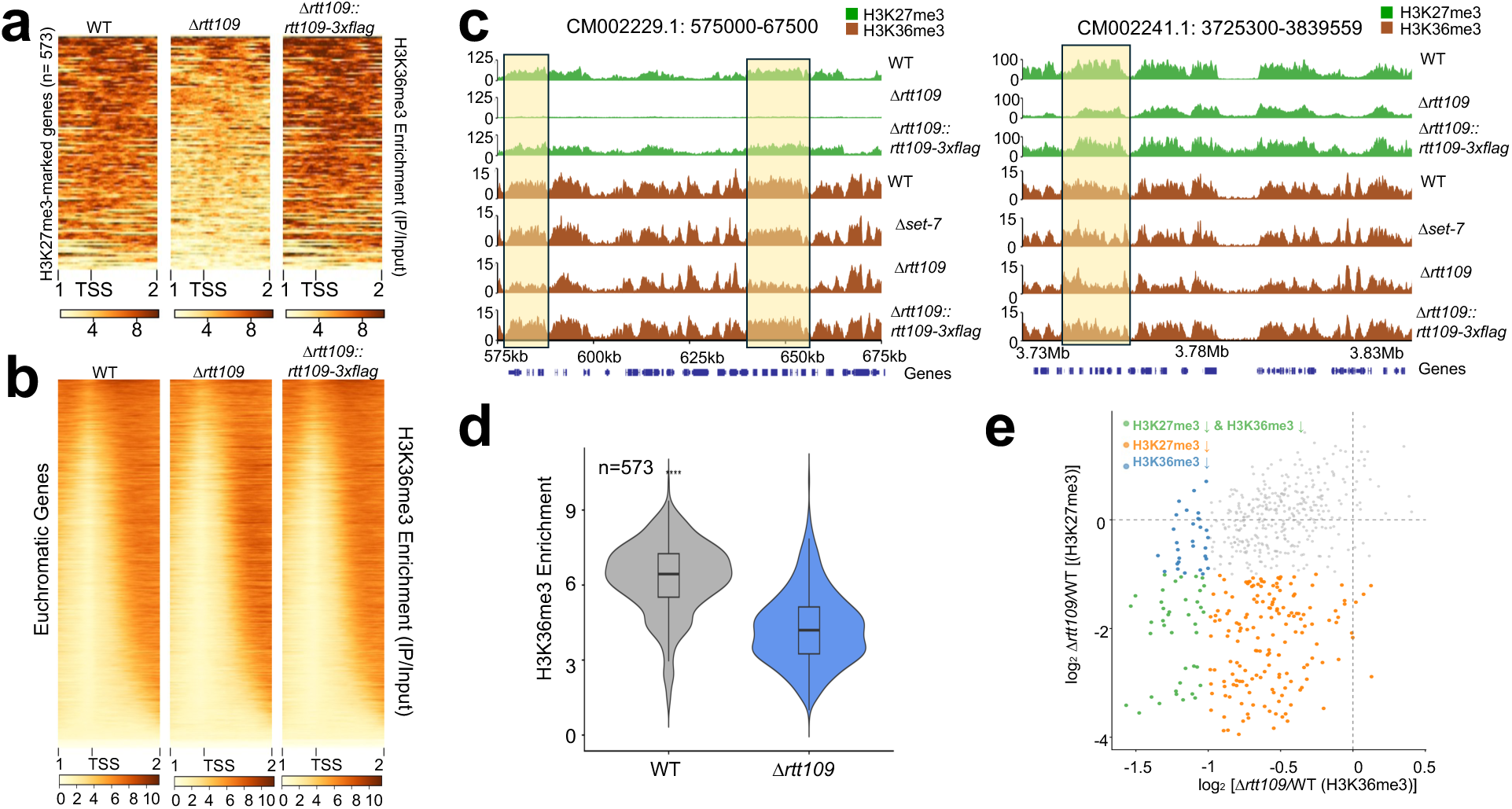
RTT109 is required for normal ASH1-dependent H3K36me3. **a.** The heatmap shows H3K36me3 enrichment across the transcriptional start site (TSS) of individual H3K27me3-marked genes (n=573), from 1 kb upstream to 2 kb downstream for the indicated strains. The heatmap is sorted from highest to lowest enrichment in the Δ*rtt109* strain. **b.** The heatmap shows H3K36me3 levels, plotted and sorted as described in panel a for euchromatic genes. **c.** The genome browser tracks show ChIP-seq enrichment of H3K27me3 (green) and H3K36me3 (orange) across a segment of LGIV (left) or LGVI (right), plotted as in fig 1c. **d.** The violin plot shows H3K36me3 enrichment across H3K27me3-marked genes (n=573) for 3 input-normalized WT and *Δrtt109* biological replicates. The box indicates the middle 50% values, the center bolded line indicates the median, the box above is the 75^th^ percentile, and the box below is the 25^th^ percentile. Asterisks indicate significance level calculated by paired Wilcoxon signed rank tests (formatted p-value < 2e-16). **e.** The scatter plot compares changes in H3K27me3 enrichment (y-axis; log_2_ [*Δrtt109/*WT]) and H3K36me3 enrichment (x-axis; log_2_ [*Δrtt109/*WT]) across H3K27me3-marked genes (n=573). Each point represents a gene (grey: log_2_ [*Δrtt109/*WT (H3K27me3)] > –1 & log_2_ [*Δrtt109/*WT (H3K36me3)] > –1; blue: log_2_ [*Δrtt109/*WT (H3K27me3)] > –1 & log_2_ [*Δrtt109/*WT (H3K36me3)] < –1; green: log_2_ [*Δrtt109/*WT (H3K27me3)] < –1 & log_2_ [*Δrtt109/*WT (H3K36me3)] < 0-1; orange: log_2_ [*Δrtt109/*WT (H3K27me3)] < –1 & log2 [*Δrtt109/*WT (H3K36me3)] > –1.

### RTT109 deficiency does not affect constitutive heterochromatin

H3K27me3 and H3K9me3 are both regarded as repressive marks but are distributed differently. In *N. crassa*, as in other eukaryotes, H3K9me3 is the hallmark of constitutive heterochromatin and is principally associated with repeat-rich domains including the centromeres. In *Schizosaccharomyces pombe*, RTT109 has been shown to be important for telomere localization at the nuclear periphery and H3K9me3 levels at constitutive heterochromatin^49–51^. To examine whether the link between RTT109 and H3K9me3 is conserved in *N. crassa*, we performed ChIP-seq on H3K9me3 on *Δrtt109*. Our results showed that loss of RTT109 does not alter H3K9me3 patterns relative to wild type, indicating constitutive heterochromatin is regulated independently of RTT109 (Supplementary Fig. 3).

### Normal H3K27me3 patterns require catalytically active RTT109 but are independent of NAF-2 and H3K56ac

In yeast, RTT109 interacts with Asf1 (Anti-silencing function 1) and Vps75p (Vacuolar Sorting Protein 75)^42^. To identify RTT109-interacting proteins in *N. crassa,* we extracted total protein from our *rtt109-3xflag* strain and performed an affinity purification using anti-FLAG resin. We then analyzed affinity-purified samples by Liquid Chromatography-Tandem Mass Spectrometry (LC-MS/MS). We identified 11 putative interacting proteins including the *Neurospora* homolog of yeast Vps75 called Nucleosome Assembly Factor 2 (NAF-2). NAF-2 was highly enriched in the RTT109 pull down, suggesting the interaction between NAF-2 and RTT109 is conserved in *N. crassa* (Supplementary Fig. 4a, Supplementary Data Tab 7). We performed a bulk pooled-protein interaction analysis using AlphaFold 3 on enriched pulled-down proteins to determine if there were other less abundant interactors^52^. Unsurprisingly, NAF-2 had the highest ipTM score (0.67) followed by NCU03300, the homolog of yeast DNA damage checkpoint component RAD24 with an ipTM of 0.63 (Supplementary Fig. 4b, Supplementary Data Tab 8). AlphaFold3 predictions identified 20 high confidence contact sites between RTT109 residues 420-480 and NAF-2 residues 290-330 (Fig. 3a, Supplementary Data Tab 9). We were unable to detect an interaction between RTT109 and ASF-1, but this is perhaps unsurprising because the Asf1-Rtt109 interaction is difficult to detect in yeast without crosslinking^53^.

**Figure 3.**
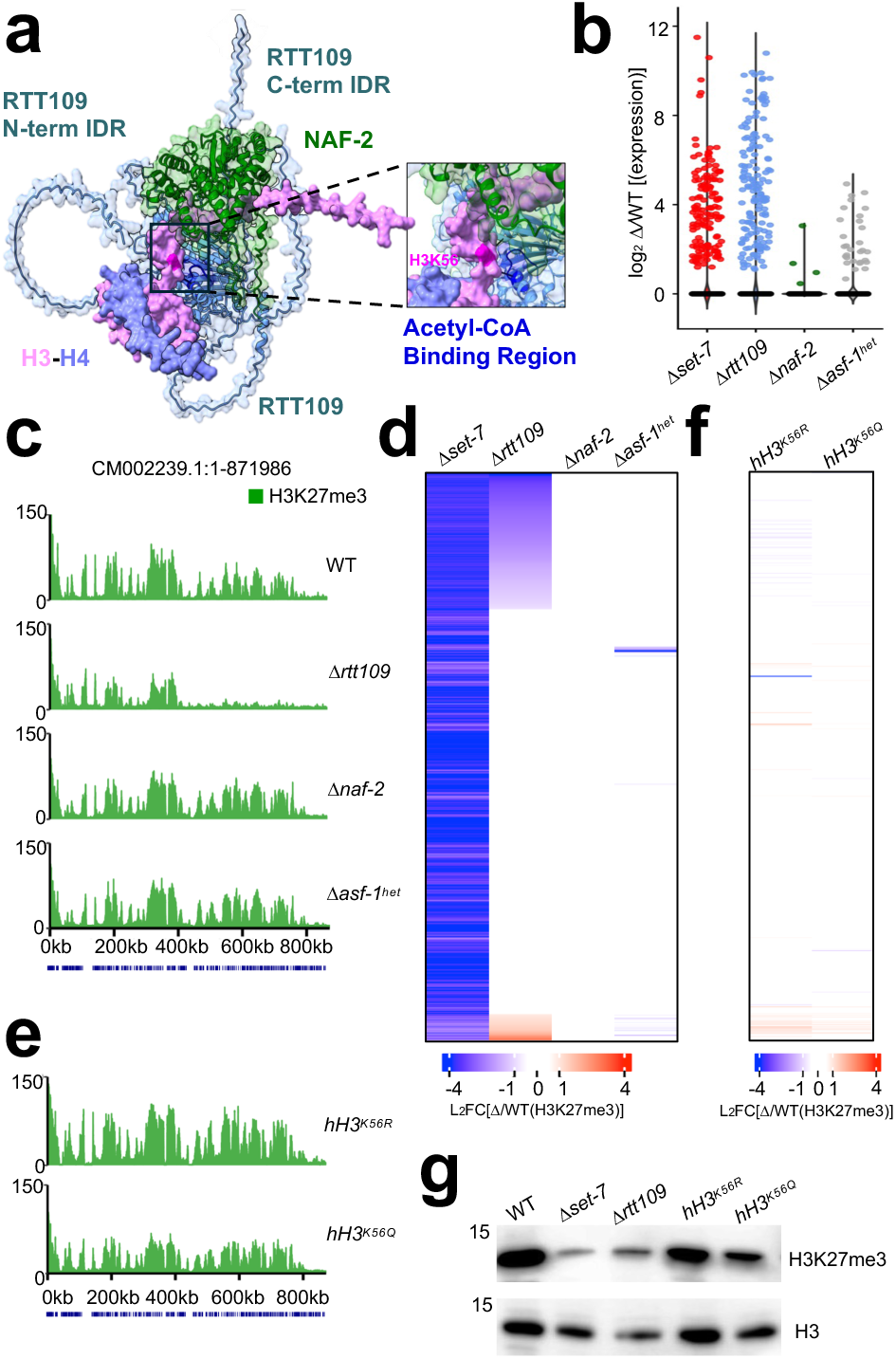
NAF-2 and H3K56 acetylation are not required for normal facultative heterochromatin. **a.** A predicted structure of the *N. crassa* RTT109-NAF-2 complex interacting with a H3-H4 heterodimer is shown. Predicted intrinsically disordered regions (IDRs) in the RTT109 C– and N-termini are labeled in blue. The inset image shows a closeup view of RTT109’s predicted acetyl-CoA binding region next to the H3K56 residue (magenta). **b.** The scatter plot shows relative expression levels (y-axis) of H3K27me3-marked genes (y-axis; n=573) for the indicated strains (x-axis; log2 [mutant/WT]). The Δ*asf-1^het^* strain contains a mixture of wild type and mutant nuclei. **c.** The genome browser tracks show normalized enrichment of H3K27me3 (ChIP-seq) for the indicated strains. **d.** The heatmap shows the changes in H3K27me3-enrichment in wild type facultative heterochromatin domains for the indicated strains. **e.** The genome browser tracks show normalized enrichment of H3K27me3 (ChIP-seq) for strains harboring the indicated H3K56 substitution alleles across the same region shown in panel c. **f.** The heatmap shows the changes in H3K27me3-enrichment in wild type facultative heterochromatin domains for the indicated strains. **g.** The western blots show levels of H3K27me3 (top) and H3 (bottom) for the indicated strains.

We asked if NAF-2 or ASF-1 was required for repression of H3K27me3-marked genes and normal H3K27me3 patterns. We performed RNA-seq on the Δ*naf-2* strain. The *N. crassa* ASF-1 gene is essential, but we also performed RNA-seq for a heterokaryotic *Δasf-1* strain (*asf-1^het^*), which had a 28% reduction in *asf-1* expression levels (Supplementary Fig. 4c). In contrast to *Δset-7* and *Δrtt109* strains, H3K27me3-marked genes were not broadly expressed in the Δ*naf-2* or *Δasf-1^het^* strains (Fig. 3b). Similarly, H3K27me3 enrichment patterns in the *Δnaf-2* and *Δasf-1^het^* strains resembled the wild type (Fig. 3c, d, Supplementary Data Tab 10).

RTT109 is the sole H3K56 acetyltransferase in fungi^42^. To determine if H3K56ac is important for facultative heterochromatin formation, we performed ChIP-seq experiments to analyze the H3K27me3 distribution in two *hH3* mutant strains in which K56 has been mutated to mimic the unacetylated or acetylated state (*hH3^K56R^* acetyl-ablative; *hH3^K56Q^* acetyl-mimetic). Neither mutant phenocopied the *Δrtt109* strain, though modest bidirectional changes in the H3K27me3 enrichment profile were observed (Fig. 3e, f; Supplementary Data Tab 10). To assess global levels of H3K27me3 in these strains, we performed an H3K27me3 western blot to detect H3K27me3 levels in crude extracts. We probed blots with antibodies against unmodified H3 as a loading control. As predicted, *Δset-7* has a strong loss of H3K27me3 compared to wild type. The remaining signal could be cross-reactivity with other modifications as previously observed^28,32^. The *Δrtt109* strain has reduced but detectable H3K27me3 levels. In contrast, H3K27me3 levels are unaffected in the *hH3^K56R^* and *hH3^K56Q^* strains (Fig. 3g). As previously reported, both H3K56 point mutants were hypersensitive to DNA-damaging agents (Supplementary Fig. 5)^45^. Taken together, our data suggest that RTT109 functions independently of NAF-2 and H3K56ac to control gene repression and repressive methylation within facultative heterochromatin domains.

We next asked if the acetyltransferase activity of RTT109 is required for steady-state H3K27me3 levels. Residues required for catalytic activity of the yeast Rtt109 ortholog are conserved in *N. crassa* RTT109^45^. We engineered two *rtt109-3xflag* alleles encoding presumed catalytically inactive proteins, *rtt109^D145A^* and RTT109^D145A-DD304-305AA^. Each of these alleles was then integrated into the *csr-1* locus of an Δ*rtt109* deletion strain. In parallel, we introduced a wild type *rtt109-3xflag* construct as a control (Supplementary Fig. 6a). H3K27me3 patterns were restored when *rtt109-3xflag* was introduced. In contrast, RTT109^D145A^ or RTT109^D145A-DD304-305AA^ failed to rescue H3K27me3 defects which was consistent with our hypothesis (Fig. 4a-c). Interestingly, the decrease in H3K27me3 was more pronounced in the strains expressing a catalytic-dead protein compared to the *Δrtt109* mutant strain. Both catalytically dead versions of RTT109 were expressed, but expression levels were lower compared to the wild type RTT109-3xFLAG protein (Supplementary Fig. 6b). Thus, we speculate that the presence of a non-functioning protein might function in a dominant negative matter, exacerbating the defect in H3K27me3.

**Figure 4.**
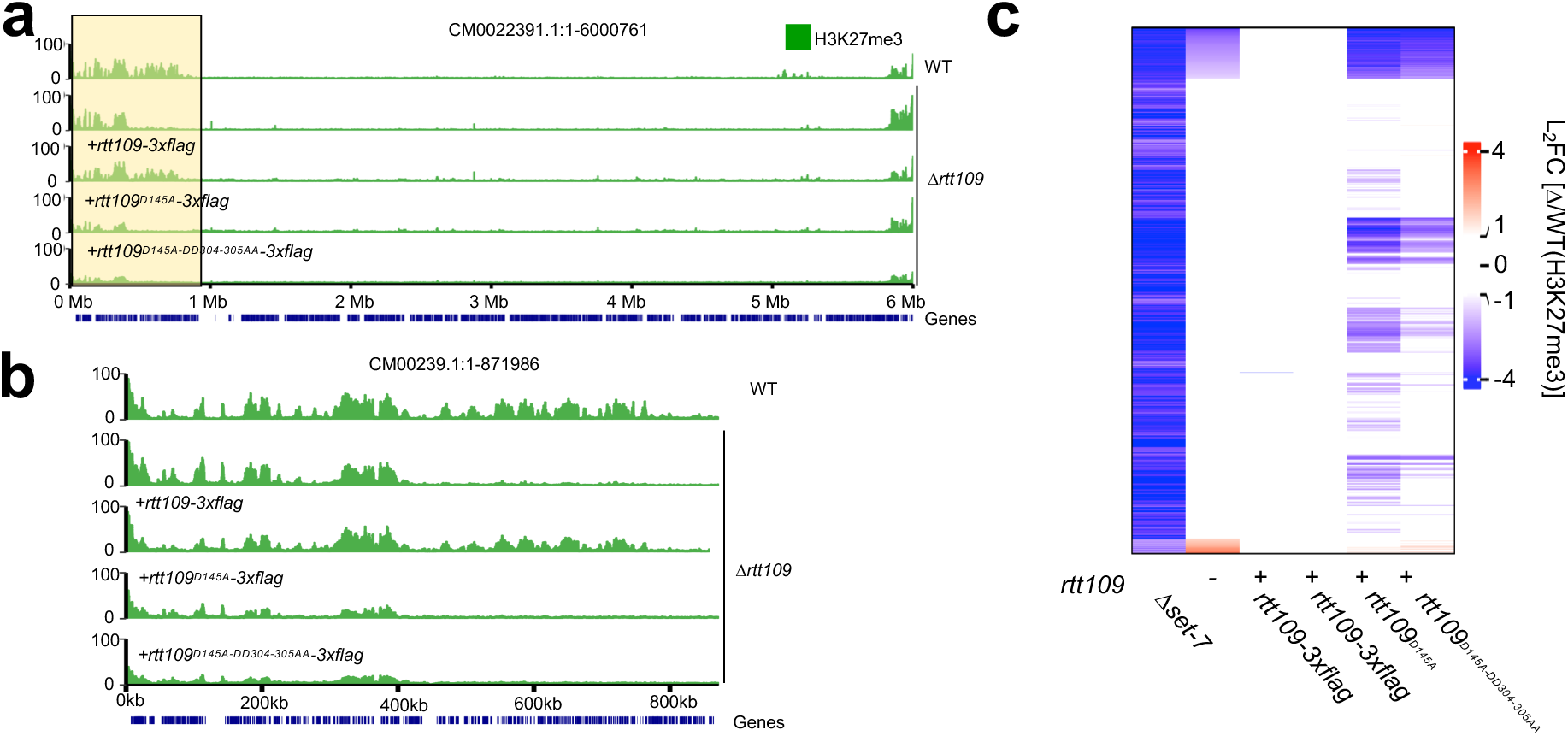
RTT109 catalytic activity is required for normal H3K27me3. **a-b**. The genome browser tracks show enrichment of H3K27me3 (ChIP-seq) across LG IV for the indicated strains. The bottom track indicates positions of annotated genes (blue). The region highlighted in the yellow box is shown in higher resolution in panel b**. c.** The heatmap shows the changes in H3K27me3-enrichment in wild type facultative heterochromatin domains for the indicated strains.

### Facultative heterochromatin decay in Δ*rtt109* occurs rapidly

Because RTT109 is known to function during replication, we reasoned that RTT109 might be important for long-term maintenance of H3K27me3 in *N. crassa*. All experiments described above were performed with the *Δrtt109* strain FGSC12340, which was generated by the Neurospora Gene Knockout Consortium and distributed by the Fungal Genetics Stock Center^54^. We do not know how many times this strain was passaged before it was obtained by our lab. If RTT109 is important for long-term stability of H3K27me3 patterns, we hypothesized that the H3K27me3 loss may be progressive, as previously observed for a strain lacking the histone deacetylase HDA-1^36^. To test this possibility, we isolated a new *Δrtt109* strain from a backcross of the *Δrtt109* FGSC12340 strain to wild type. We then sequentially passaged this backcrossed strain *(Δrtt109* P0) and performed ChIP-seq to assay H3K27me3 after each passage (Supplementary Fig. 7a). Based on the average number of spores produced in a slant, we estimate that each passage represents approximately ∼25-30 mitotic divisions^55^. In the Δ*rtt109* P0 isolate, H3K27me3 enrichment was reduced compared to wildtype but was significantly higher than in the parental Δ*rtt109* FGSC12340 strain (Fig. 5a). We observed progressive loss of H3K27me3 in sequentially passaged Δ*rtt109* strains (P1-3; Fig. 5b). H3K27me3 ChIP-qPCR on *ncu04869* confirmed these results (Supplementary Fig. 7b, c). We also passaged the *hH3^K56R^* strain and found no changes in the H3K27me3 enrichment pattern between the original strain and a strain passaged three times, consistent with a role for RTT109 that is independent of H3K56 acetylation (Supplementary Fig. 7d). We assayed H3K36me3 across passaged *Δrtt109* strains via ChIP-seq to determine whether H3K36me3 loss is also progressive. The newly isolated P0 strain already displayed the striking loss we saw in the parental strain from Figure 2 (Supplementary Fig. 7e). This suggests that, in contrast to H3K27me3, loss of RTT109 leads to a more rapid reduction of H3K36me3. Taken together, our results suggest that RTT109 is critical for H3K36me3 in facultative heterochromatin and required for the long-term stability of H3K27me3 patterns.

**Figure 5.**
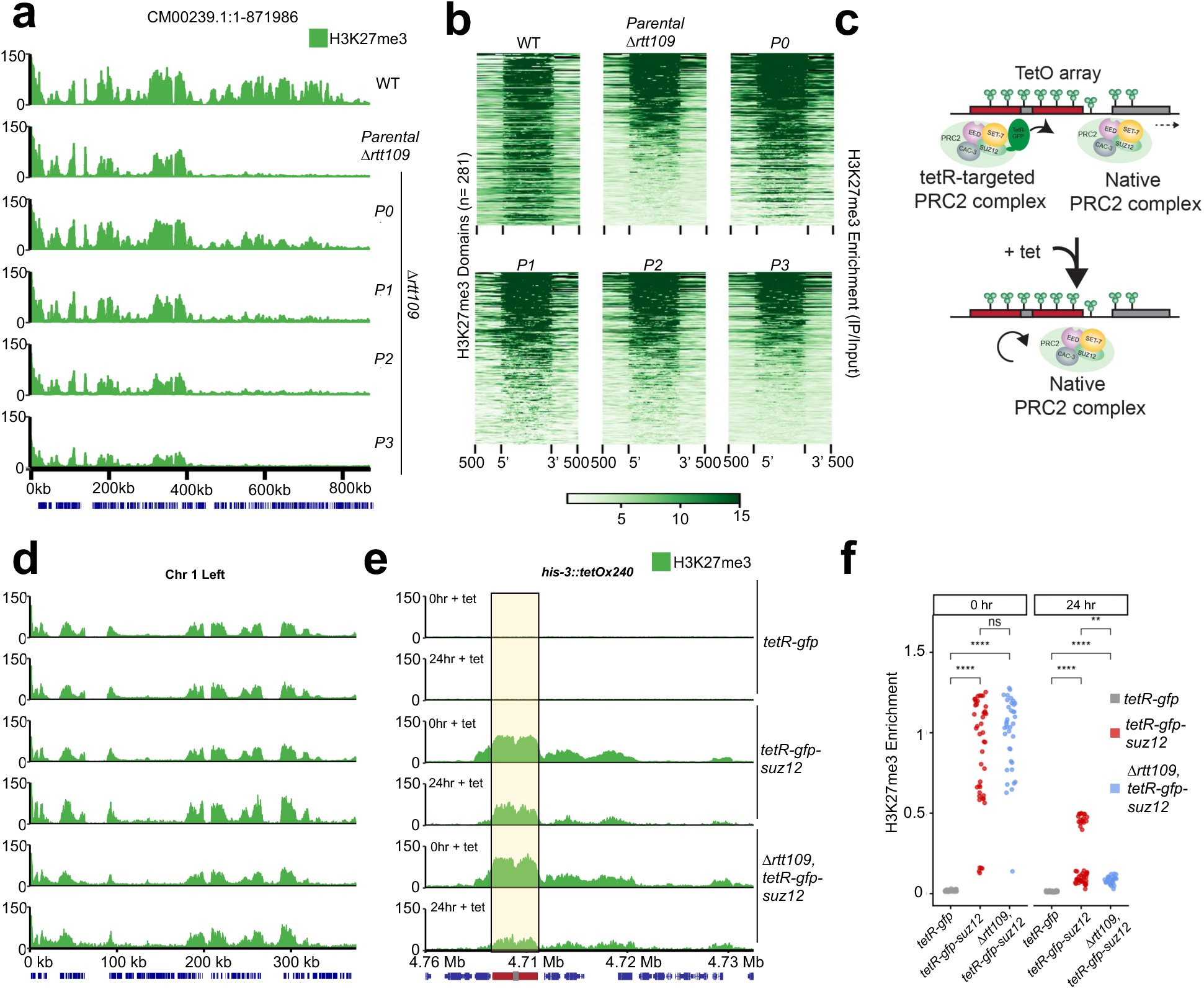
RTT109 is important for H3K27me3 maintenance. **a.** The genome browser tracks show enrichment of H3K27me3 (ChIP-seq) across a section of LG IV for the indicated strains. The bottom track shows gene positions (blue). Chromosomal coordinates are shown on the x-axis in kb. The bottom track shows positions of annotated genes. Tracks are shown for wild type (WT), the parental *Δrtt109* strain (P0) or a new *Δrtt109* strain isolated from a backcross (P1) and sequentially passaged (P2-3). **b.** The heatmap shows H3K27me3 enrichment (ChIP-seq) across wild type H3K27me3 domains (n=281) with ±500 bp flanking regions on either side (rows) for the strains in panel a. Domains are sorted from highest to lowest enrichment in the parental *Δrtt109* strain. **c.** The schematic diagram illustrates the experimental approach used to reversibly target PRC2 to a tetO array integrated at the *N. crassa his-3* locus. Tetracycline is added to block binding of the tetR^OFF^-GFP-SUZ12 construct to the *tetO* array. **d-e.** The genome browser tracks show H3K27me3 enrichment within a native facultative heterochromatin region near the left arm of LG I (d) or within the *tetO* array (e) for the indicated strains in before addition of tetracycline (0hr + tet) or after growth in tetracycline for 24 hours (24hr + tet). The bottom track indicates positions of annotated genes (blue) or the *tetO* array (red). **f.** The scatter plots show normalized H3K27me3 enrichment for the indicated strains at 0 and 24 hours. H3K27me3 enrichment values are normalized to the 95th percentile value in the time-matched tetR^OFF^-GFP control. Asterisks indicate significance level calculated by paired Wilcoxon signed rank tests (**** = formatted p-value < 0.0001, ** = formatted p-value < 0.01, ns = not significant).

### RTT109 is required for mitotic stability of H3K27me3

Our passaging experiments suggest that RTT109 may be important for maintenance of H3K27me3 patterns over multiple mitotic divisions. To test this possibility, we developed a synthetic system to reversibly target PRC2 to a specific locus and we used this system to separately assay the establishment, spreading, and maintenance of H3K27me3 *in vivo*. We constructed a *tetR^OFF^-gfp-suz12* fusion construct and introduced this into a strain harboring an array of *tet* operator sequences^56^ (*tetO*). In this system, *tetR^OFF^* constitutively tethers the tetR-SUZ12 fusion to the *tetO* array enabling targeted H3K27me3 deposition, while the addition of tetracycline dissociates the fusion protein from the locus (Fig. 5c). Addition of this construct did not affect endogenous H3K27me3 (Fig. 5d; Supplementary Fig. 8a, b, c). In minimal medium lacking tetracycline, the tetR^OFF^-GFP-SUZ12 established a new H3K27me3-enriched domain at the *tetO* array (Fig. 5e). H3K27me3 was highly enriched across the entire *tetO* array and appeared to spread asymmetrically into adjacent euchromatic genes. As expected, the control strain expressing tetR^OFF^-GFP, which lacks any PRC2 subunit, showed no enrichment of H3K27me3 at *tetO* (Fig. 5e; Supplementary Fig. 8a). Next, we asked if the native PRC2 complex would maintain H3K27me3 at *tetO* after disassociation of tetR^OFF^-GFP-SUZ12. In wild type *N. crassa*, H3K27me3-enrichment persists for at least 24 hours following addition of tetracycline to the media (Fig. 5e). We estimate that 10-12 rounds of nuclear division occur during this

24-hour period^55^. Together, these results demonstrate that H3K27me3 directed to *tetO* can be epigenetically maintained across mitosis without continuous PRC2 recruitment.

To determine if RTT109 is important for establishment of H3K27me3 by tetR^OFF^-GFP-SUZ12 or for maintenance of H3K27me3 in the presence of tetracycline, we deleted *rtt109* in the *tetO; tetR^OFF^-gfp-suz12* background. When this Δ*rtt109* strain was grown in media lacking tetracycline, H3K27me3 was highly enriched at the *tetO* array and spread into adjacent genes similar to the wild type strain (Fig. 5e). In contrast, we observed significantly reduced H3K27me3-levels when these Δ*rtt109* cells were grown in the presence of tetracycline (Fig. 5d-f, Supplementary Fig. 8d, Supplementary Data Tab 11). Taken together, these data suggest that RTT109 is required for long-term maintenance of H3K27me3 in the absence of continuous PRC2 recruitment.

## DISCUSSION

In plants, animals, and some fungi, PRC2 forms stably repressed chromatin domains to control cell type-specific gene expression. Assembly and maintenance of PRC2-repressed chromatin are thought to involve three mechanistically distinct but functionally coordinated processes: (1) *de novo* recruitment of PRC2 to establish H3K27me3, (2) spreading of H3K27me3 along the chromatin fiber, and (3) epigenetic inheritance mechanisms that reassemble repressed chromatin after each nuclear division^2,6,13^. PRC1 was lost early in the fungal lineage, raising the possibility that H3K27me3 maintenance does not occur in fungi or involves PRC1-independent mechanisms^25^. Here, we show that H3K27me3 is stably maintained over multiple mitotic divisions in *N. crassa* and this epigenomic stability depends on RTT109, a replication-coupled acetyltransferase.

How cells preserve epigenetic information through the disruptive process of DNA replication remains a central question in chromatin biology. Early investigations in *S. pombe* revealed heritability of H3K9me3 relies on a “read-write” feedback loop where the methyltransferase (Clr4) recognizes the mark it creates^57,58^. Stable maintenance of H3K9me3 depends on pre-existing modifications, hypoacetylated histones, dose-dependent H3 ubiquitylation and recruitment of multiple chromatin regulators^59–62^. Faithful inheritance of these heterochromatin domains is further sustained by balanced parental histone segregation and replisome factors such as FACT, CAF-1, and Mrc1^63–65^. Our finding that RTT109 is required for stable maintenance of H3K27me3 adds to the growing body of evidence that key replication components are critical for preserving epigenetic memory^17,20,21,24,63,64^.

Originally discovered in budding yeast, Rtt109 works with histone chaperones Vps75 and Asf1 to acetylate newly synthesized H3K56 and residues in the H3 tail (K9 and K27)^42^. RTT109-dependent acetylation of lysines in the H3 tail is thought to modulate replication speed to ensure proper nucleosome assembly^66,67^, while H3K56ac significantly increases H3-H4 binding affinity for the CAF-1 histone deposition complex^68^. Once incorporated into the genome, H3K56ac has been shown to stimulate activity of the ATP-dependent chromatin remodeler Isw1 (Imitation Switch 1) to facilitate proper nucleosome spacing and chromatin maturation^69^. Thus, studies in yeast have established RTT109 as an important regulator of RCNA and chromatin maturation. Our findings reveal that RTT109 plays an additional, mechanistically distinct role in maintaining facultative heterochromatin in filamentous fungi.

In *N. crassa,* RTT109-deficient strains inappropriately express genes in facultative heterochromatin and exhibit region-specific losses of two repressive histone modifications, H3K27me3 and ASH1-dependent H3K36me. Perhaps surprisingly, we show here that H3K56 acetylation is not required for normal structure and function of facultative heterochromatin. This conclusion is supported by transcriptomic and H3K27me3 ChIP-seq data for H3K56 substitution mutants and *Δnaf-2*. Although *hH3^K^*^56^ point mutants did not phenocopy Δ*rtt109* strains with respect to H3K27me3 patterns, these strains were defective for DNA repair as previously reported^45^. In contrast, RTT109’s acetyltransferase activity is required for normal H3K27me3 patterns. We note that protein levels of catalytically inactive RTT109 were reduced compared to wild type, and therefore we cannot eliminate the possibility that H3K27me3 loss reflects reduced RTT109 levels rather than loss of catalytic activity. However, this seems unlikely given that strains expressing RTT109^D145A^ or RTT109^D145A-DD304-305AA^ exhibit H3K27me3 losses that are similar or greater to the *Δrtt109* strain.

We propose two possible models for RTT109’s function in maintaining facultative heterochromatin (Fig. 6). It is possible that RTT109 acetylates a histone residue other than H3K56. Although acetylation is generally associated with transcriptional activation, it is possible that transient histone acetylation increases the accessibility of the H3 tail to facilitate methylation by PRC2 or ASH1. Indeed, in the Δ*hda-1* mutant of *N. crassa,* hyperacetylation of centromeres was associated with increased DNA methylation and aberrant H3K27me3 deposition at these sites, consistent with the idea that acetylation can increase activity of repressive enzymes^36,70^. Biochemically, Rtt109-Vps75 can also acetylate H3K9 and H3K27. However, our *Δnaf-2* mutant strain did not exhibit a defect in H3K27me3, suggesting that acetylation of these residues is also not required for maintenance of facultative heterochromatin.

**Figure 6.**
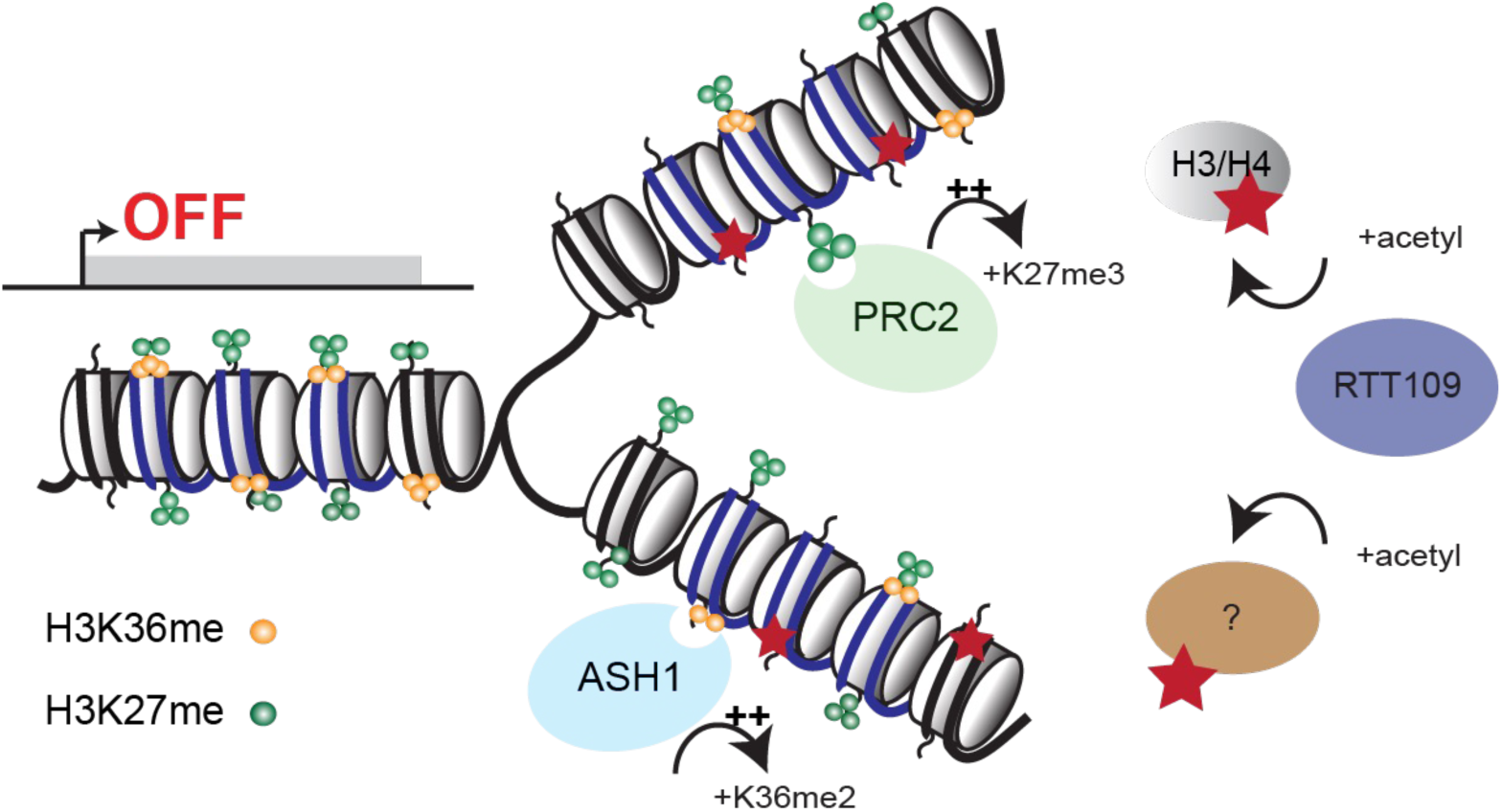
RTT109 controls epigenetic maintenance of H3K27me3 through H3K56-independent acetylation. The schematic diagram illustrates possible mechanisms by which RTT109 regulates epigenetic inheritance of the repressed chromatin state.

Alternatively, RTT109 may acetylate a non-histone protein. RTT109 is a structural homolog of mammalian p300/CBP^71^, which has both histone and non-histone substrates^72^. An acetyl proteomics study in *N. crassa* identified 940 proteins and 1940 acetylated lysine residues^73^. Several known or potential regulators of facultative heterochromatin were found to be acetylated including the chromatin remodeler ISW, the histone deacetylase RPD3, a putative ASH1-interacting protein, and others. H3K36me3 is rapidly lost in early passages of the *Δrtt109* mutant, raising the possibility that RTT109 regulates ASH1-dependent H3K36 methyltransferase activity. PRC2 is another potential substrate of RTT109. Acetylation of mammalian EZH2 alters its stability^74^. However, PRC2 subunits were not identified in the *N. crassa* acetylome^73^. Further investigation to identify the relevant RTT109 substrate is an important goal for future studies. We note that our RNA-seq analysis of additional *N. crassa* acetyltransferase knockout strains confirmed that RTT109 is the sole histone acetyltransferase required for gene repression in facultative heterochromatin, suggesting that this is a specific function of RTT109.

The functional importance of RTT109 as a key chromatin regulator is evident across filamentous fungi. In *Aspergillus flavus,* loss of RTT109 is associated with decreased secondary metabolite biosynthesis, growth defects and poor DNA damage repair^75^. Unsurprisingly, RTT109 is required for virulence in fungal pathogens of plants and animals such as *Candida albicans*, *Beauveria bassiana*, and *Aspergillus fumigatus*^76–80^. The discovery that RTT109 is important for mitotic stability of gene expression states may provide a potential mechanistic explanation for some of these functions. Because RTT109 is fungal-specific, it presents a potential therapeutic target for treating infections by targeting epigenetic plasticity that underlies pathogenicity and environmental adaptation. More broadly, this work demonstrates that PRC1-independent mechanisms support mitotic inheritance of H3K27me3-repressed chromatin in fungi and highlight the importance of replisome-associated proteins in maintaining the epigenome.

## Materials and Methods

### Strain construction and growth

Strains used in this study are listed in Supplementary Table S1. Strains were grown at 32°C in Vogel’s Minimal Medium (VMM) + 1.5% sucrose for ChIP-seq, RNA-seq, DNA, and protein isolation^81^. Liquid cultures were shaken at 180 rpm. Crosses were performed on Synthetic Crossing (SC) medium in the dark at room temperature and individual spores were isolated as described^81^.

*DNA Transformation*: To create a targeted complementation strain, we designed a *rtt109-3xflag-TrpCp-bar* knock-in construct using primers from Supplementary Table S2. The construct was cloned by Gibson assembly into the pCSN43 plasmid backbone using the NEBuilder HiFi DNA Assembly Master Mix (NEB, Cat. # E2621L). The construct (600 ng) was transformed by electroporation into a *Δrtt109::hph, Δmus-51:hph* strain as described^82^. Individual colonies were isolated 4-5 days after transformation on VMM containing 2% sorbose, 0.1% glucose, 0.1% fructose (FGS) and BASTA salts, 3% Proline, and 200 µg/mL BASTA. Transformants were confirmed by testing for complementation of the MMS sensitivity phenotype, PCR, and Western blot using an anti-FLAG antibody (Sigma Aldrich, Cat. # F1804-200UG). Catalytic RTT109 mutants were designed based on conserved residues identified by homology from *S. cerevisiae* and cloned by Gibson assembly into a *csr-1* targeting vector pEAG249G^83^ (NEBuilder HiFi DNA Assembly Master Mix; Cat. # E2621L) and maintained in *E. coli*. Transformants were selected on VMM containing 5 µg/ml cyclosporin A and maintained on slants containing 1 mL 1xVMM + 1.5% sucrose + 10 µg/ml cyclosporin A. All plasmids were confirmed by Oxford Nanopore Sequencing (Plasmidsaurus).

*Reporter Strain Generation*: For generating the *ncu06889::bar^OFF^* reporter strain, we designed a *ncu06889* replacement construct in which the coding sequence was replaced with a basta resistance cassette gene (*bar,* driven by the native *ncu06889* promoter). We transformed the DNA into the *Δset-7, Δmus-52* strain, selected for a basta-resistant transformant, and backcrossed the strain to wild type to isolate the *ncu06889::bar^OFF^* reporter in a *set-7+, Δmus-52+* background. We then crossed the reporter to the Δ*rtt109* (S534) and Δ*eed* (S603) strains to test for heterochromatic gene silencing defects.

*tetR-PRC2 strain*s: A *csr-1* integration plasmid containing *pDim5-tetR-gfp-suz12* was constructed by Gibson Assembly (NEBuilder HiFi DNA Assembly Master Mix; Cat. # E2621L). The *suz12* coding sequence was amplified by PCR using genomic DNA as a template and cloned into pEAG247C (Supplementary Tables 1,2). The *his-3::tetO; Δcsr-1::tetR-gfp-suz12* strain (S872) was made by transforming strain S543, a gift from Eugene Gladyshev. Primers were used to amplify *Δrtt109::hph* from FGSC12340 and this fragment was transformed into strain S872 (*csr-1::pdim5-tetR-gfp-suz12, his-3::tetO-240x*). A single transformant was passaged 3 times to isolate a homokaryon. To disassociate the tetR^OFF^-GFP-SUZ12 fusion from chromatin, strains were grown in the absence of tetracycline for 12 hours and exposed to 2.5 µg/mL tetracycline for 24 hours, the 0-hour control was never exposed to tetracycline before being harvested for ChIP-seq.

*Spot tests:* Spot tests for the reporter strain were done on VMM plates containing FGS, 200 µg/mL Hygromycin B Gold (Invivogen, Cat. # ant-hg-1), or 200 µg/mL BASTA. Conidia (asexual spores) were serially diluted and spotted onto plates containing VMM, VMM + 200 µg/mL Hygromycin B, and VMM + 0.010% (w/v) methyl methanesulfonate (Sigma Aldrich, Cat. # 129925-5g). Plates were imaged after a 48-hour incubation at 32°C.

### Chip-seq, RNA-seq, and molecular biology

*Chromatin immunoprecipitation (ChIP):* Conidia were inoculated in 5 mL of VMM + 1.5% sucrose and grown for ∼18 hours. ChIP was performed as described previously^84^. In brief, mycelia were harvested using vacuum filtration and were washed once in 1xPBS prior to cross-linking in 1xPBS + 1% formaldehyde for 10 minutes. The reaction was quenched by addition of 125mM glycine and subjected to an additional wash with 1xPBS. Mycelial tissue was resuspended in 600 µl of ChiP lysis buffer (50mM HEPES, pH 7.5, 140mM NaCl, 1mM EDTA, 1% Triton X-100, 0.1% sodium deoxycholate, one tablet Roche cOmplete mini EDTA-free Protease Inhibitor Cocktail (Sigma Aldrich, Cat. # 11836170001)). Chromatin was sheared by sonication with the QSONICA Misonix S-4000 ultrasonic processor and Diagenode Bioruptor UCD-200. Lysates were centrifuged at 13,000 rpm for five minutes at 4°C. Supplementary Table S3 lists antibodies used in this study. Immunoprecipitation was performed by incubating the lysate with antibody and Protein A/G beads (20 µl) (Santa Cruz, Cat. # sc-2003) overnight. Beads were washed twice with 1 mL ChIP lysis buffer, once with 500mM NaCl ChIP Wash buffer, once with 50mM LiCl ChIP Wash buffer, and finally with TE Buffer. Bound chromatin was eluted in ChIP TES Buffer (50mM Tris pH 8.0, 10mM EDTA, 10% SDS) at 65°C for 10 minutes and de-crosslinked overnight at 65°C. The DNA was treated with RNase A for two hours at 50°C, then with proteinase K for two hours at 50°C and extracted via ActivMotif Chromatin IP DNA Purification Kit (Cat. #. 58002) or Zymo ChIP DNA Clean & Concentrator Kit (Cat. # D5205) and eluted in 32 µl diH_2_O. Samples were then prepared for Illumina sequencing or ChIP-qPCR.

*ChIP library preparation:* Libraries were constructed as described previously^84,85^. In brief, the NEBNext Ultra II End Repair/dA-tailing Module (Cat. # E7546S), NEBNext Ultra II Ligation Module (Cat. # E7546) were used to clean and A-tail DNA prior to Illumina adapter ligation. The ligation products were amplified to generate dual-indexed libraries using NEBNext Ultra II Q5 Hot Start HiFi PCR Master Mix (Cat. # M0543S). Size selection with Sera-Mag SpeedBeads (Cytiva, Cat. # 65152105050250) was performed after the adapter ligation and PCR steps. Beads are suspended in a solution of 20mM PEG 8000, 1mM NaCl, 10mM Tris-HCl, 1mM EDTA at 4°C prior to use.

*ChIP-qPCR:* 3-4 biological replicates of ChIP input samples were diluted 1:50 and IP samples 1:10 before proceeding with qPCR. Reactions were set up using Biorad Universal SYBR Green Mastermix (Cat. #4309155) based on the manufacturer’s specification. ChIP-qPCR was performed using ThermoFisher QuantStudio™ 3 System Instrument 272323132 with experimental set up Comparative-Ct-SYBR (ΔΔCт) where *gh-3-6* is the control (Supplementary Table 1). Two-tailed student’s t-tests were used to determine significant H3K27me3 enrichment.

*FLAG Co-Immunoprecipitation and Protein Complex Purification*: 7-10 day old conidial cultures were harvested and used to inoculate 1 liter of 1xVMM + 1.5% sucrose and grown with typical growth rates. Mycelia were harvested by vacuum filtration, washed with 1xPBS, flash-frozen, and ground in liquid nitrogen. Mycelial powder was incubated in protein complex purification (PCP) buffer (150 mM HEPES pH 7.5, 1mM EDTA pH 8.0, 250 mM KCl, 10% Glycerol, 1mM DTT) at 4°C until thawed on a rotator. Samples were subjected to centrifugation at 2000 rpm for 5 minutes and the supernatant was transferred to a new tube before addition of 200 µl anti-FLAG resin (Sigma Aldrich, Cat. #A36801). The lysate was incubated with anti-FLAG resin overnight at 4°C. Resin-bound protein was washed six times in PCP prior to elution using 200 µg/mL 3×FLAG peptide (Sigma Aldrich, Cat. # A2220-5ml). Proteins were concentrated by trichloroacetic acid precipitation. Eluted proteins were run 5 minutes on a 4–12% SDS–PAGE gel (Biorad, Cat. # 3450124), stained with SYPRO Ruby (ThermoFisher, Cat. # S12000), excised as a single band, digested with trypsin, and analyzed on a ThermoFisher LTQ Orbitrap Elite at the UGA Proteomics and Mass Spectrometry Facility. False discovery rate is represented as the Peptide Spectrum Match score as determined by Mascot.

*Western Blotting*: Proteins were extracted using 50mM HEPES, 150mM NaCl, 0.1% SDS, 1mM EDTA, 1mM PMSF buffer and quantified using Pierce BCA Assay Kit (ThermoFisher, Cat. #23225). 10-20µg total protein was boiled in 2x Sample Buffer (Biorad, Cat. # 1610737EDU) for 5 minutes at 95°C and loaded onto SDS-PAGE gels. Proteins were transferred using the Trans-blot Turbo System (Biorad, Cat. #1704150) using the preset HMW setting and stained with Ponceau (ThermoScientific, Cat. #A40000279) before immunoblotting. All blocking steps were performed for 1 hour at room temperature (RT), primary antibody incubations occurred at 4°C overnight, and secondary antibody were for 35 minutes at RT. Membranes were rinsed 3 times for 5 minutes with 1xTBST in-between incubations.

FLAG blots were blocked with 5% milk (1xTBST) and incubated with primary 1:5000 anti-flag mouse (Sigma, Cat. # F1804-200UG) in 5% milk (1xTBST), followed by secondary 1:25000 goat anti-mouse (Life Technologies, Cat. # A11004). For H3K27me3 blots, membranes were blocked with 5% BSA (1xTBST), incubated with primary 1:2000 anti-H3K27me3 mouse (Diagenode, Cat. # NBP2-59295), followed by secondary 1:25000 goat anti-mouse antibody (ThermoFisher, Cat # 31450) 5% BSA (1xTBST). H3 blots were blocked with 5% BSA (1xTBST), incubated with primary 1:2000 anti-Histone H3 rabbit (Active Motif, Cat. # 61799), followed by secondary 1:25000 goat anti-rabbit IgG antibody (Active Motif, Cat # 15015) 5% BSA (1xTBST). SuperSignal™ West Femto Maximum Sensitivity Substrate (ThermoFisher, Cat. # 34094) was used according to manufacturer’s specifications to develop the blot prior to imaging. Uncropped membranes and quantification by FIJI are available in Source Data.

## Data Analysis

*RNA-seq Analyses:* For RNA-seq data, short reads (< 25 bp) and adaptor sequences were removed using TrimGalore (version 0.6.10-GCCcore-12.3.0)^86^. Paired-end trimmed reads were aligned to the current *N. crassa* NC12 genome (NCBI:GCA_000182925.2) using STAR/2.7.11b-GCC-13.3.0 with an intron max of 10kb^87^. Sorted BAM files were indexed using SAMtools/1.21-GCC-13.3.0^88^. Reads that overlapped coding sequences were counted using featureCounts (Subread/2.0.6-GCC-12.3.0) and grouped by gene name^89^. Counts were used in differential gene expression analysis (R package DeSeq2) for 3 biological replicates per strain^90^. Genes with fewer than 6 reads across all samples were excluded, and non-significant (adjusted p-value > 0.01) genes were set to log_2_ [*Δ*/WT (expression)] = 0.

*ChIP-seq Analyses*: Short reads (< 25 bp) and adaptor sequences were removed using TrimGalore prior to alignment to *N. crassa* NC12 GCA_000182925.2 using BWA (version 0.7.17)^86,91^. Files were sorted and indexed using SAMtools^88^. Genome-wide coverage was calculated in 50 bp bins using DeepTools (version 3.3.1) and normalized by base per million (BPM) to generate bigwig files. Replicate BAM files were merged using SAMtools prior to background correction with bamCompare (DeepTools) to generate log_2_[ChIP/Input]^92^. MACS3 (version 3.0.1) was used to identify H3K27me3-marked genes and peaks in WT and *Δrtt109* (compared to input controls) using paired-end BAM files from 3 biological replicates per strain, a genome size of 41.0 Mb, broad peak calling, FDR = 0.1, minimum domain length of 800 bp, and a maximum gap of 500 bp^47^. Differential H3K27me3 enrichment was quantified using CSAW in 300 bp windows filtered at > log_2_(2.5) over global background (n= ∼7k), normalized to their respective inputs, and to comparison against WT^93^. If CSAW calculated FDR > 0.05, then log_2_ values were set to 0. To quantify H3K27me3 enrichment at the *tetO* array, H3K27me3 enrichment values at *tetO* were calculated in 300bp windows using CSAW. Each window log_2_(enrichment/input) was normalized to the 95th percentile of genome-wide H3K27me3 signal (excluding the *tetO* and the mating-type locus). Statistical analyses were performed as pairwise comparisons using a Wilcoxon rank sum-test.

*Data Visualization:* Input-normalized bigwigs were used for genome browser track by R package rtracklayer^94^ and DeepTools for heatmaps, line plots, and Spearman correlation matrix (Figure S5)^92^. Computematrix was used for mapping H3K27me3 domains or H3K9me3 domains with the parameters, –scale regions –-skipZeros –b 500 –a 500 –-sortUsingSamples Δrtt109 –-sortRegions descend and excluding zero-coverage bins. plotHeatmap then sorted genes by the sum of H3K27me3. enrichment (green) across H3K27me3 domains using Δrtt109. For gene level analyses, H3K27me3 or H3K36me3 enrichment was calculated over gene bodies extended by 1 kb upstream and 2 kb downstream of the transcription start site (TSS) based on *Neurospora crassa* gene annotations (GCA_000182925.2_NC12_genomic).

*AlphaFold3*: AlphaFold3 predictions between RTT109 and candidate proteins were run on UGA Sapelo2 cluster according to guidelines made by GACRC (https://wiki.gacrc.uga.edu/wiki/AlphaFold3-Sapelo2)^52^. ipTM, average pTM scores and standard deviation were pulled from output.json files. Size correction was not required based on the ipTM and amino acid length across the pooled protein-protein prediction interactions. NAF-2 and RTT109 were modeled in UCSF ChimeraX with a H3-H4 heterodimer for figure 3a.

## Data Availability

All sequencing data associated with this study are available through public databases at the National Center for Biotechnology Information (NCBI). RNA-seq data were downloaded from the NCBI short read archive (accession numbers listed in Supplementary Data Tab 12). ChIP-seq data generated for this study are available from the Gene Expression Omnibus under series record #GSE330894. Custom scripts used to analyze or visualize high throughput sequencing data are publicly available on github: https://github.com/ry00555/RTT109_SourceCode

## Acknowledgements

Support for this work comes from the NIH/NIGMS Grant #R35GM152134. We thank Daphnee Dubreuil for technical contributions to this work. We thank Drs. Joel Steyer and Quanita Choudhury for critical reading of the manuscript. We thank Drs. Qun He and Eugene Gladyshev for sharing *N. crassa* strains.

## Notes

### Competing Interest Statement

The authors have declared no competing interest.

