## SupplementaryFiguresAndTables for "Epigenetic maintenance of PRC2-repressed chromatin requires RTT109 but not H3K56 acetylation"

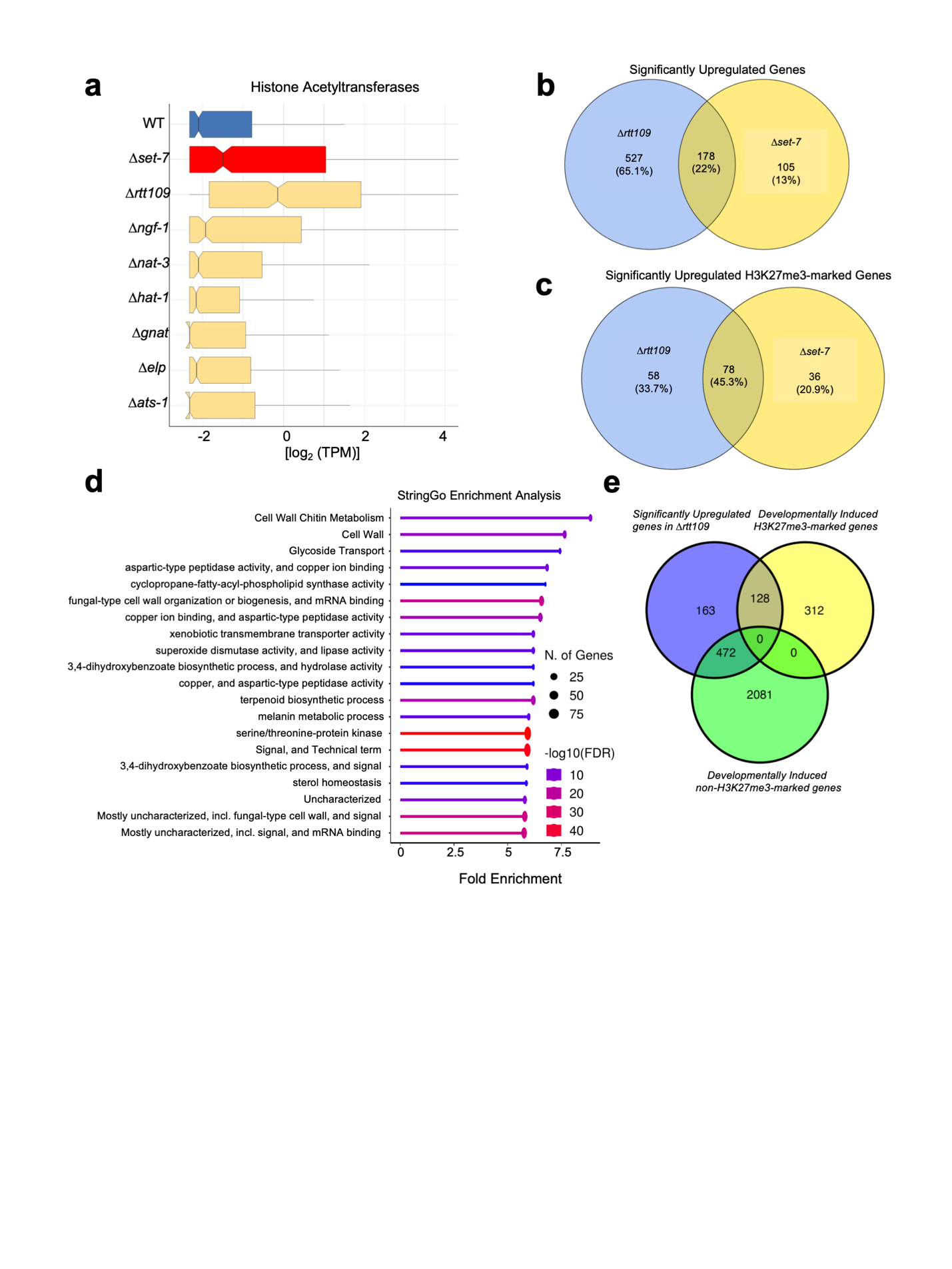
Supplemental Figure 1: A comparative analysis between *∆rtt109* and other mutants reveal RTT109 is required for repression of genes in facultative heterochromatin domains. **a.** The boxplots show the average level of expression for all H3K27me3-marked genes (n= 573) for the indicated acetyltransferase mutant strains, with the x-axis values corresponding to log_2_-transformed transcripts per million (TPM) values. Lines represent the median expression value, and the notches indicate the 95% confidence interval. **b.** The venn diagram shows number of overlapping significantly upregulated genes between *∆rtt109* and *∆set-7.* **c.** The venn diagram shows number of overlapping or upregulated H3K27me3-marked genes between *∆rtt109* and *∆set-7*. **d.** A dot plot summarizing StringGo Enrichment Analysis on upregulated genes in *∆rtt109*. **e.** The venn diagram shows number of overlapping significantly upregulated genes in *∆rtt109* and developmentally induced genes marked with and without H3K27me3.

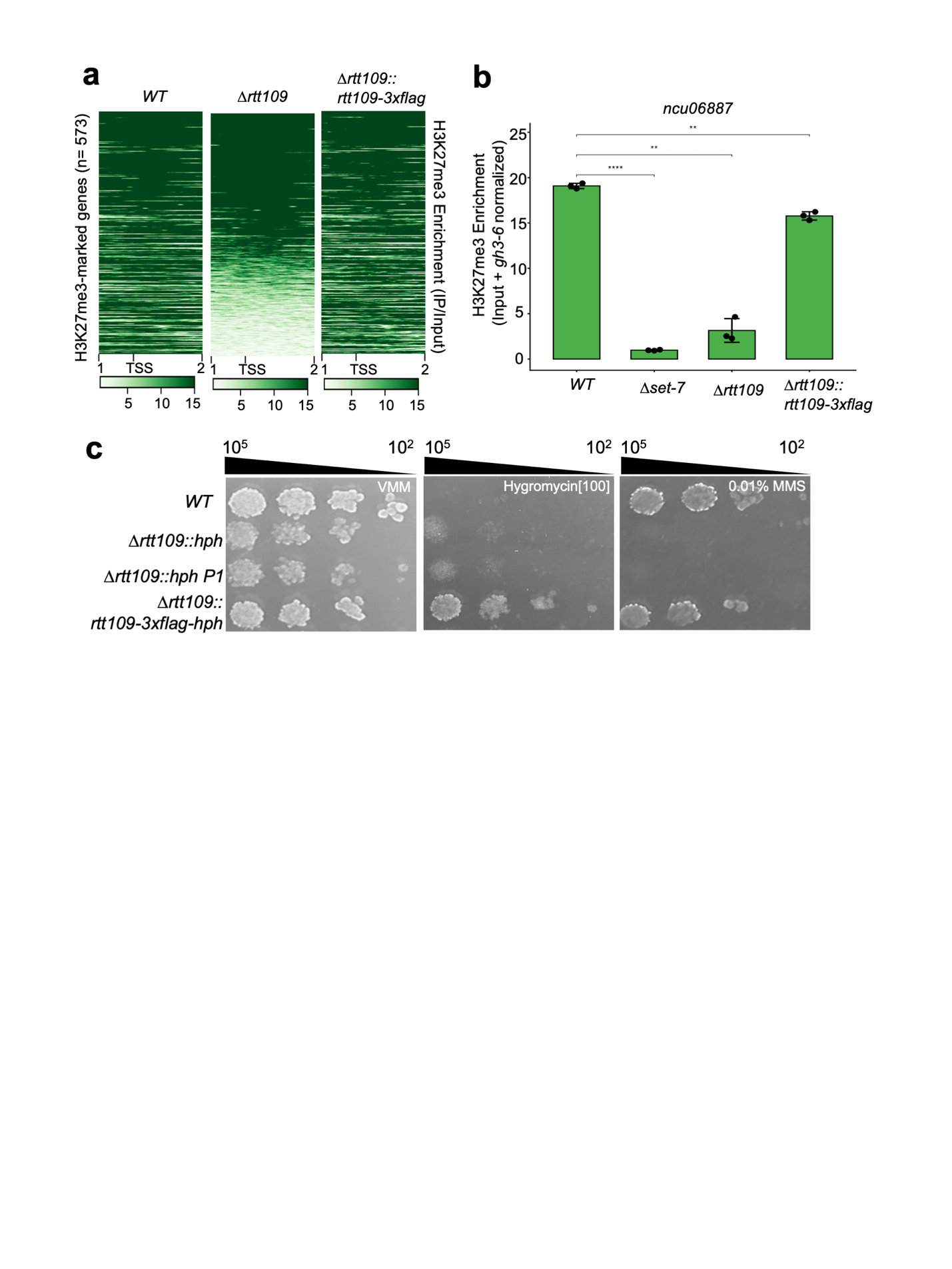

Supplemental Figure 2: *∆rtt109* causes changes in H3K27me3 patterns and can be rescued by complementation. **a.** The heatmap shows enrichment along individual H3K27me3-marked genes (n=573) with 1 kb upstream and 2 kb downstream of the transcriptional start site (TSS). **b.** The bar chart depicts ChIP-qPCR data of H3K27me3 enrichment levels over *ncu06887* for 3 biological replicates per strain*.* Error bars represent standard deviation Asterisks indicate significance level calculated by paired Wilcoxon signed rank tests (**** = formatted p-value < 0.0001, ** = formatted p-value < 0.01, ns = not significant). **c.** The spot test shows *∆rtt109* DNA-Damage sensitivity phenotype is restored in the complemented strain. Media types and dilution series is at the top indicates how many cells were plated.

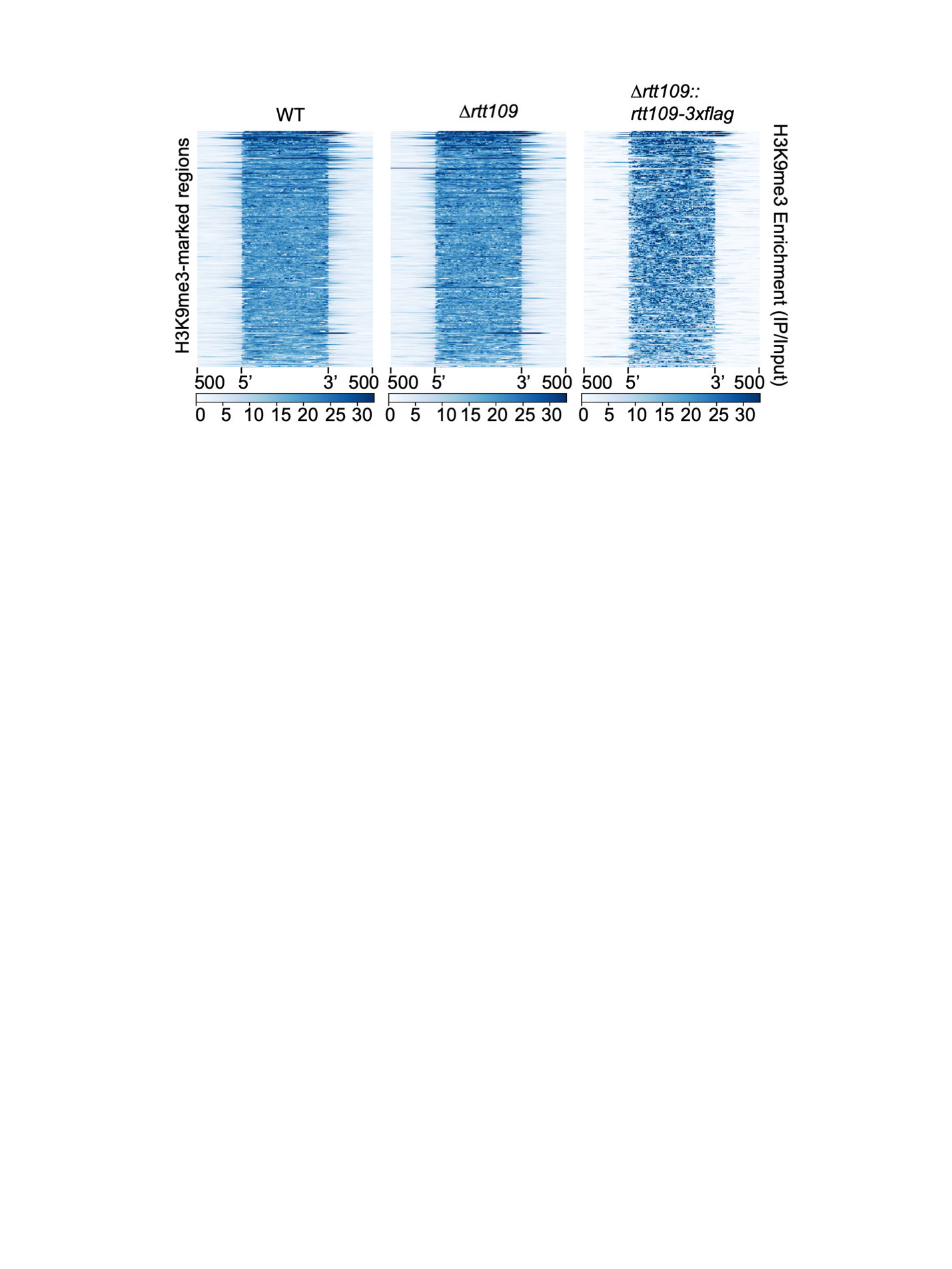

Supplemental Figure 3. Loss of RTT109 does not cause defects in H3K9me3 in constitutive heterochromatin domains. The heatmap contains H3K9me3 enrichment across all WT constitutive heterochromatin regions ±500 bp for the indicated strains.

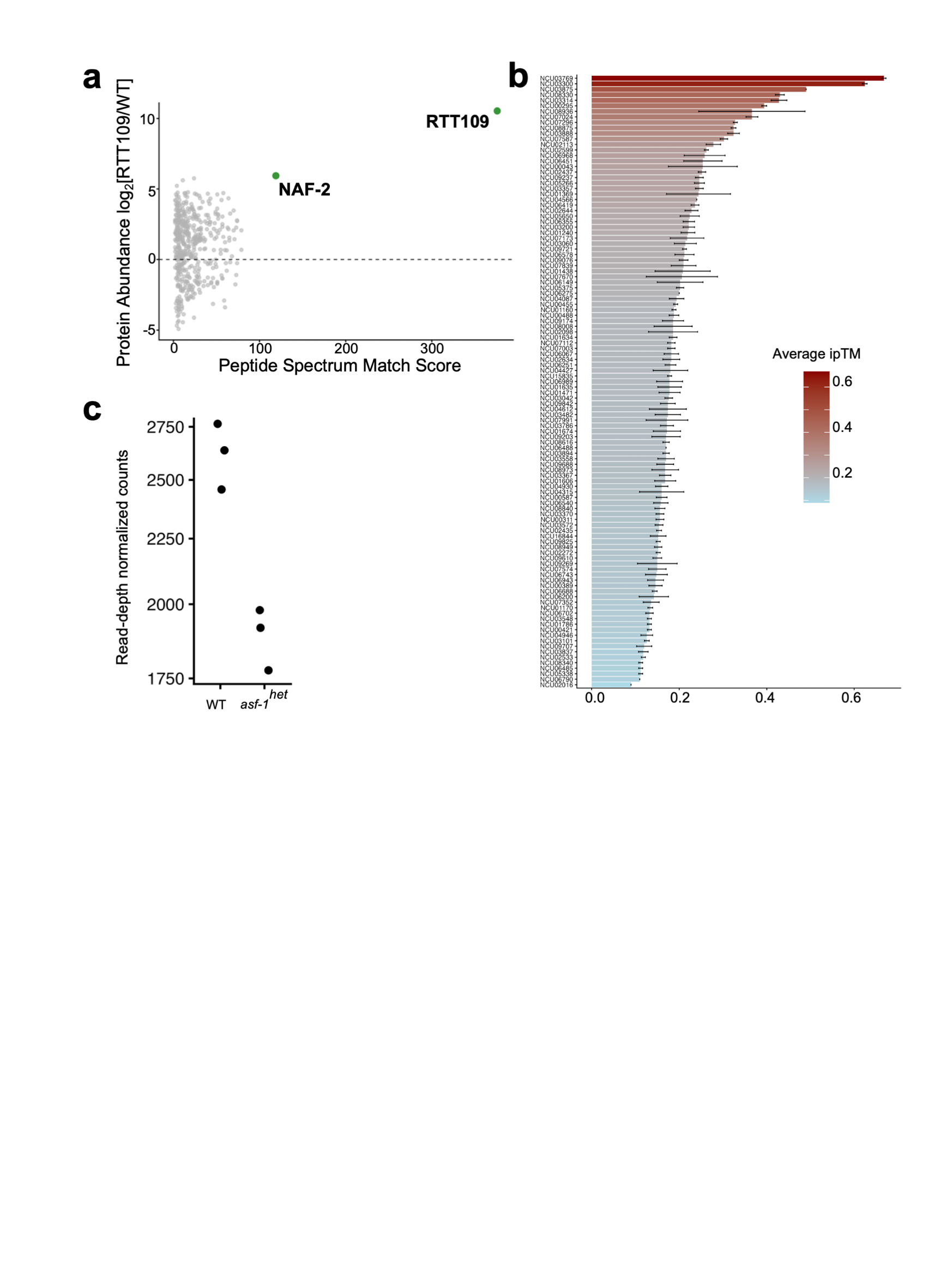

Supplemental Figure 4. **a.** The volcano plot shows peptide enrichment scores (PSM) versus log_2_ fold change of relative enrichment (RTT109/WT) from mass spectrometry analysis. Protein enrichment was calculated by normalizing the number of unique peptides relative to global protein abundance derived from PaxDB. Proteins passing a combined significance threshold (log_2_FoldChange > 5 and SumPEPScore above the 40^th^ percentile) are highlighted in green. Labeled proteins represent high-confidence interactors or known chromatin regulators enriched in the RTT109 complex. **b.** The bar chart shows average ipTM scores and standard deviation from AlphaFold3 predictions between RTT109 and proteins identified from mass spectrometry. **c.** The dot plot depicts normalized read counts across the *asf-1* locus in WT and in *asf-1^het^* strain.

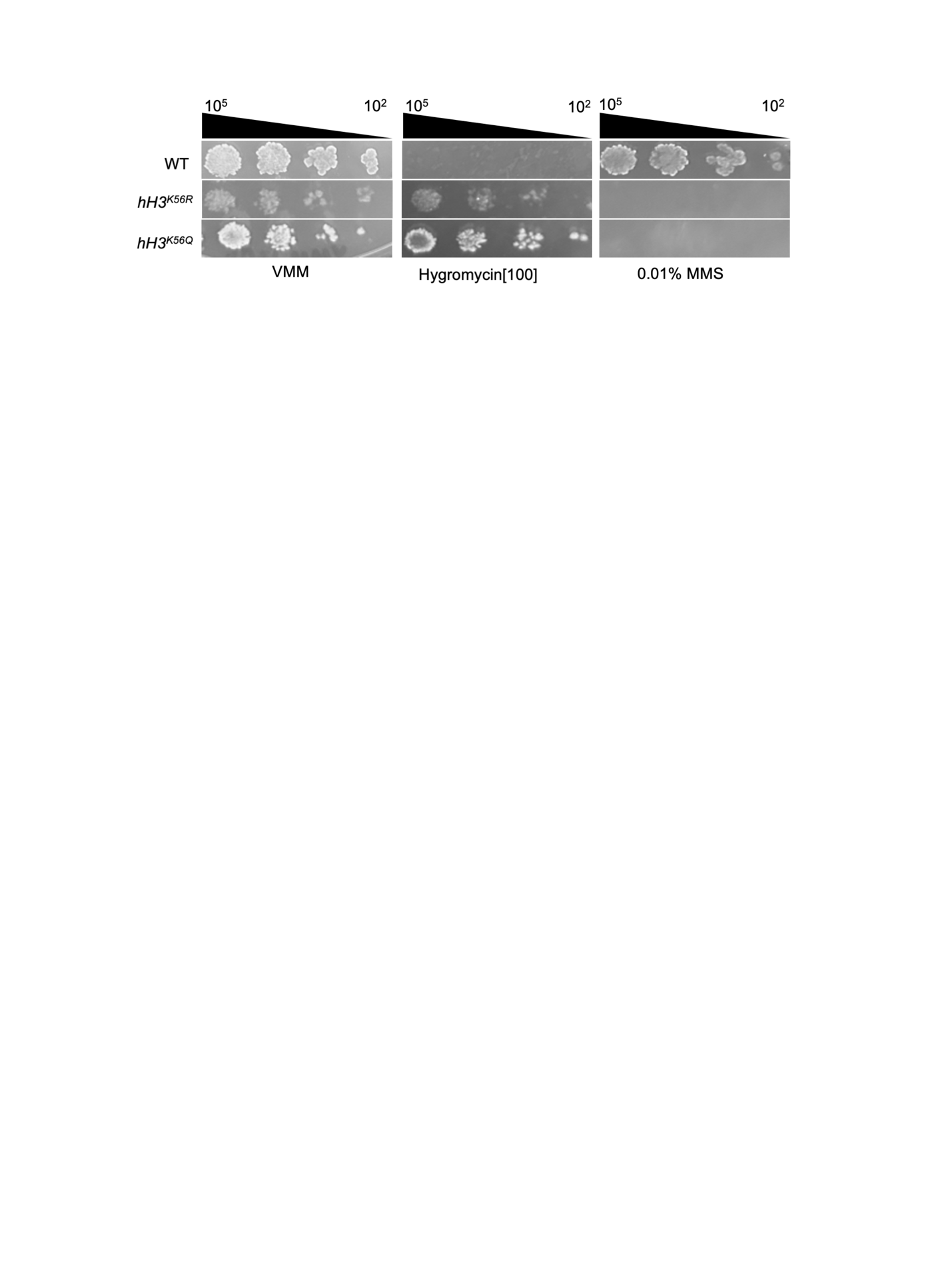

Supplemental Figure 5: H3K56 point mutants are phenocopy DNA Damage sensitivity. The spot test assays methyl methanesulfonate (MMS) sensitivity for the indicated strains.

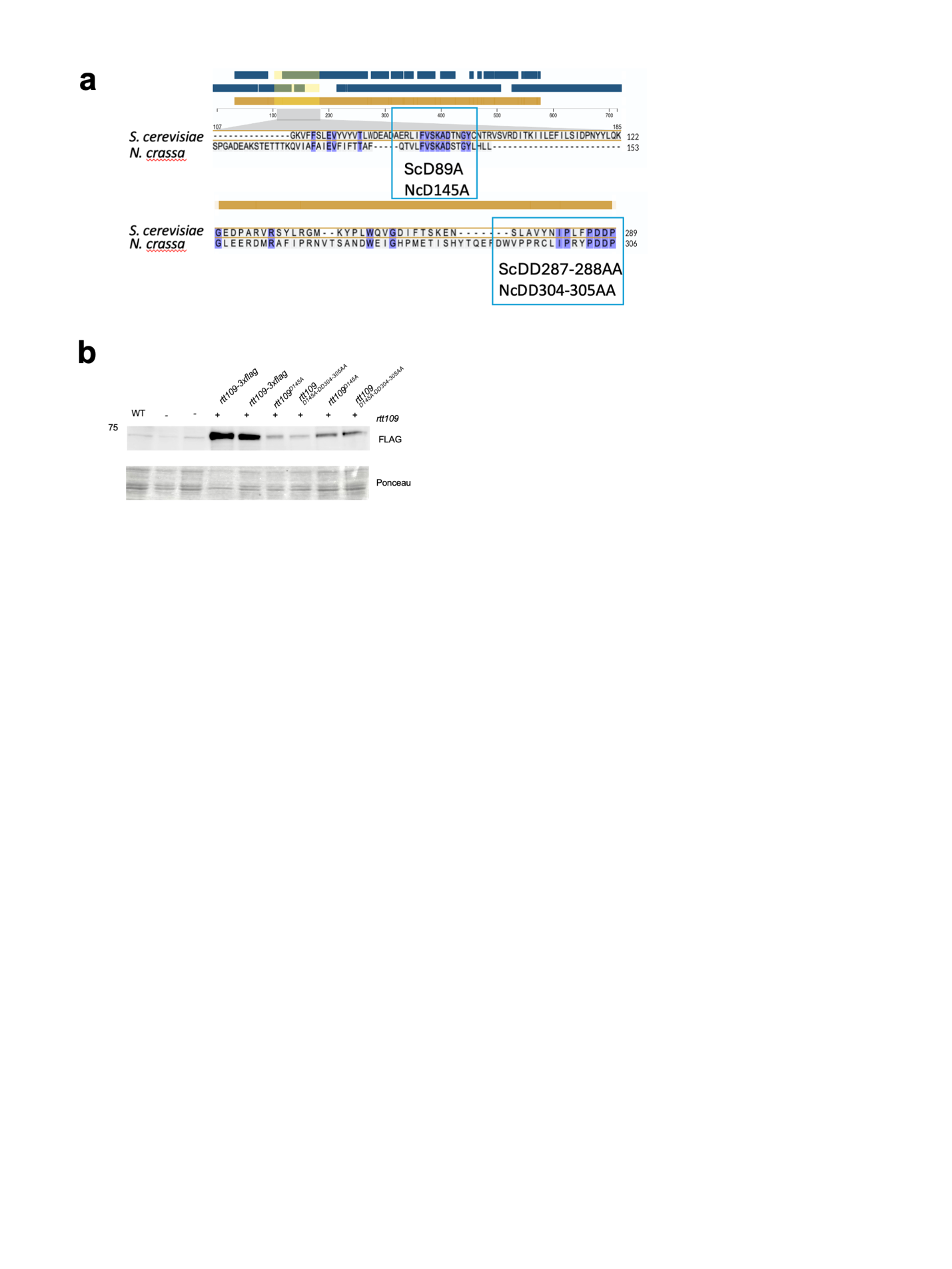

Supplemental Figure 6: Systematic mutagenesis of *rtt109* to create a catalytic dead mutant. **a.** UniProt Alignment of Neurospora crassa NCU09825 (RTT109) amino acid sequence aligned to *Saccharomyces cerevisiae* Rtt109. Blue boxes highlight the conserved aspartate residues required for catalytic activity. **b.** FLAG Western blot for indicated strains. Lanes are labeled using the following key: 1)rtt109::rtt109::3xflag, 2) ∆rttt109::hph, csr-1::rtt109-3x-flag, 3,5) ∆rtt109::hph, csr-1::rtt109^D145A^-3xflag 4,6) ∆rtt109::hph, csr-1::rtt109^D145A-DD304-305AA^-3xflag. An equal amount of crude protein (10 µg) was used for WT, ∆rtt109, and Samples 1-4. Samples 5 and 6 had twice the amount of protein (20 µg) loaded to highlight the lower amount of RTT109 protein in the rtt109D145A and rtt109^D145A-DD304-305AA^ mutants.

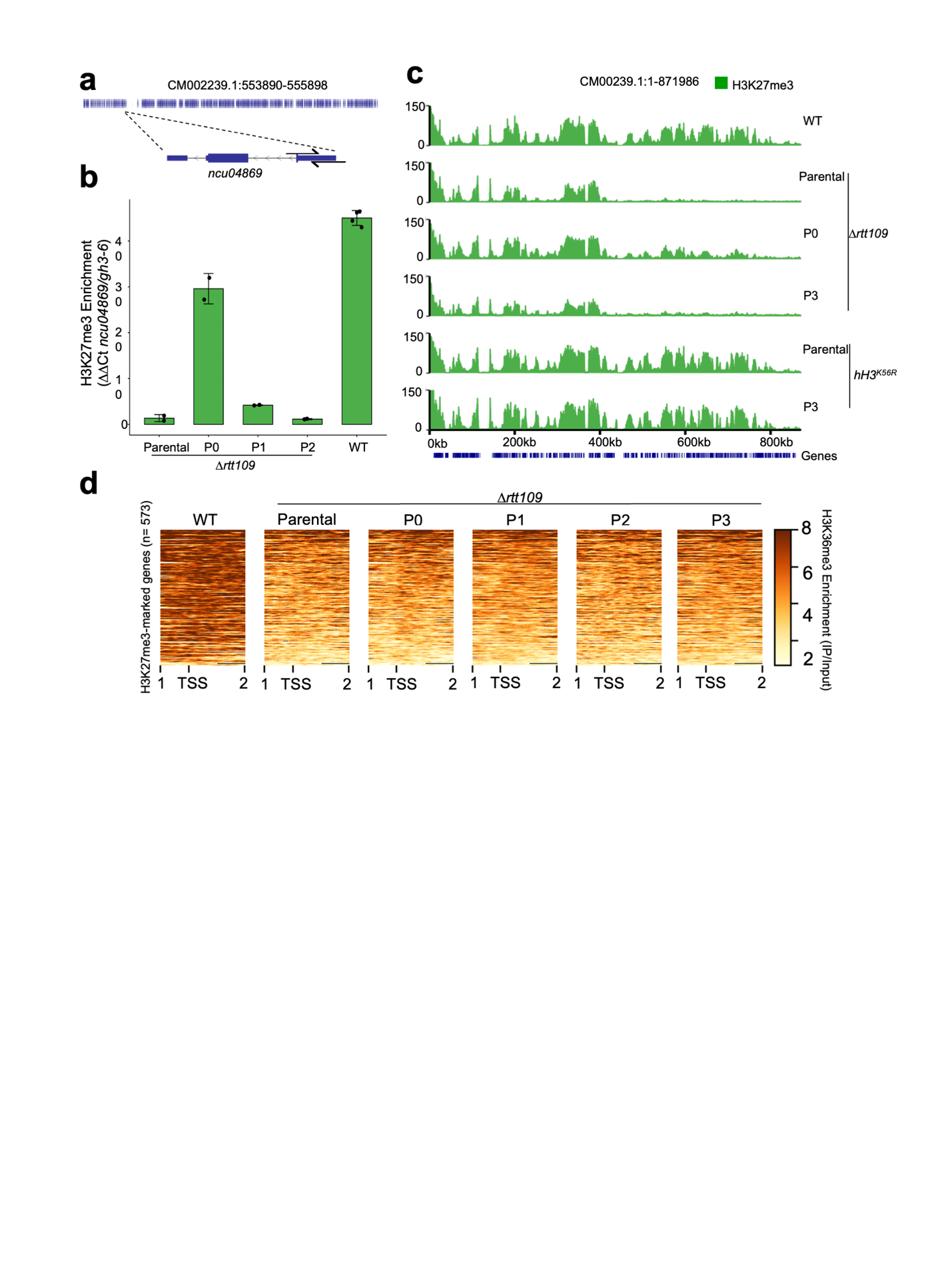

Supplemental Figure 7: **a.** Schematic of *ncu04869*, an H3K27me3-marked gene found in the region where H3K27me3 is lost in *∆rtt109*. Black arrows indicate where the ChIP-qPCR primers used in Supplemental Figure 6B bind to. **b.** A bar plot summarizing ChIP-qPCR experiment to assess H3K27me3 enrichment at the *ncu04869* locus in the indicated strains from Figure 5A. Ct values from biological replicates were grouped by strain, and the mean Ct value is shown as a bar for each strain. Error bars represent the standard deviation across replicates. Individual data points correspond to Ct values from each replicate and are overlaid as dots. Lower Ct values indicate higher enrichment of H3K27me3 at the locus. **c.** The genome browser tracks show ChIP-seq enrichment of H3K27me3 (green) across a specified fragment in LG IV. **d.** Heatmap shows H3K36me3 enrichment along individual H3K27me3-marked genes (n=573) with 1 kb upstream and 2 kb downstream of the transcriptional start site (TSS) for the indicated strains. Genes (rows) were sorted by the sum of H3K36me3 enrichment (brown) in across H3K27me3-marked genes in parental *∆rtt109*.

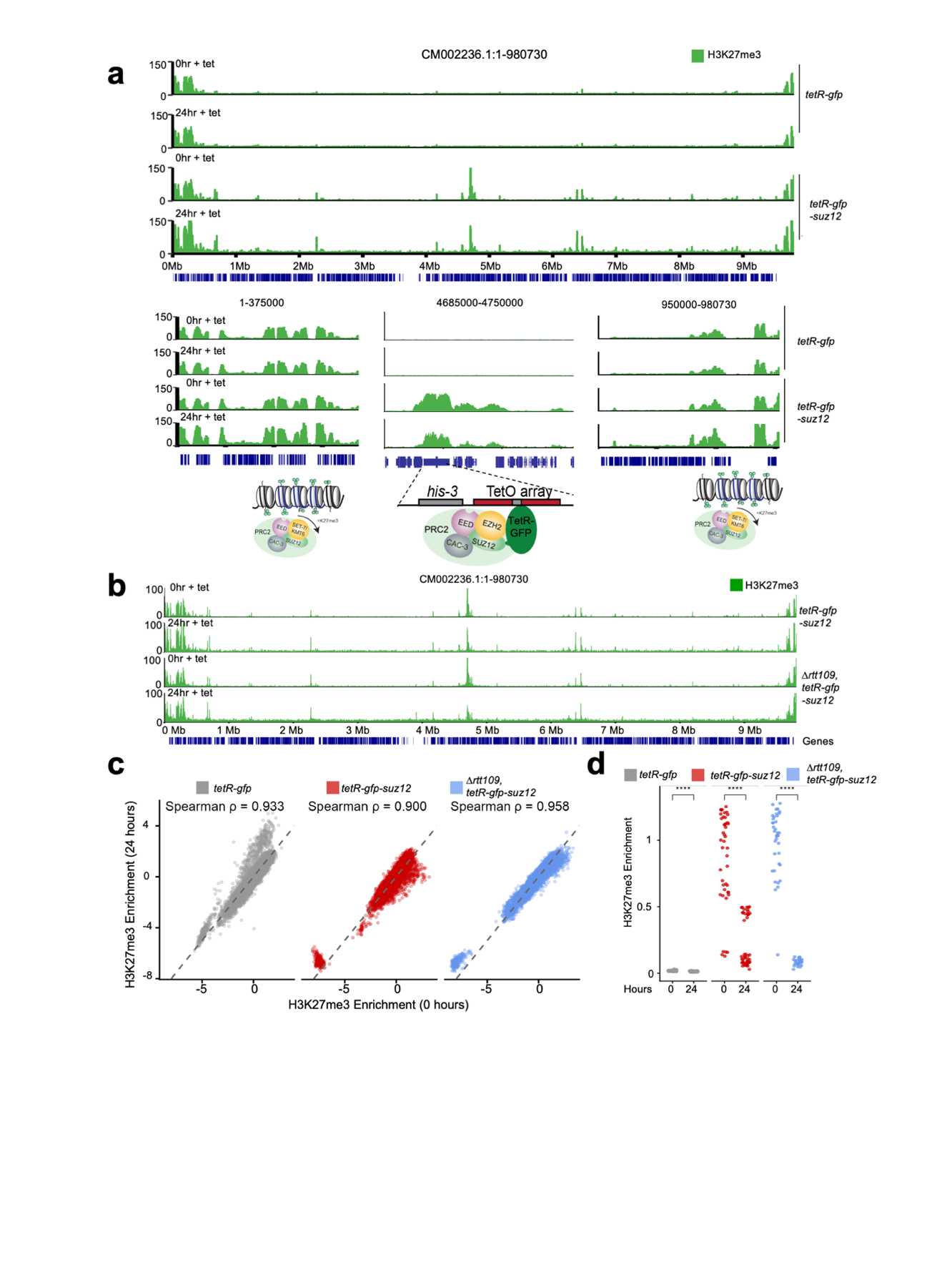

Supplemental Figure 8. **a.** A genome browser shot of chromosome 1 showing the distribution of H3K27me3 (green) enrichment of the end of chromosome 1 (left), tetO array at the *his-3* locus (center), and end of chromosome 1 (right). The number of hours represents the number of hours in which the cultures were exposed to tetracycline. Below the tracks are the genes in blue. **b.** The genome browser tracks show H3K27me3 enrichment for the indicated strains harboring across the same region shown in panel a. **c.** Spearman correlation on H3K27me3 enrichment found in the genome excluding tetO and mating type locus for the indicated strains. **d.** The dot plots show H3K27me3 enrichment at tetO sites for the indicated strains at 0 and 24 hours. Asterisks indicate significance level calculated by paired Wilcoxon signed rank tests (**** = formatted p-value < 0.0001, ** = formatted p-value < 0.01, ns = not significant).

| Supplemental Table 1. List of primers used in this study. | | |
| --- | --- | --- |
| Primer name | Sequence | Purpose |
| twist fp | CAATCCGCCCTCACTACAACCG | Used to amplify catalytic-dead rtt109 constructs to put into pBM61 |
| twist rp | TCCCTCATCGACGCCAGAGTAG | Used to amplify catalytic-dead rtt109 constructs to put into pBM61 |
| RY59-Gibson-Inverse-Primer-2 | GTGGACGGCTAATGGGGTCTGAATGCTAAA | Used to amplify pBM61 backbone for csr-1 integrations |
| RY99-pBM61GibsonInverseExtended | AGCTCCAATTCGCCCTATAGTGAGTCGTATTAAGG | Used to amplify pBM61 backbone for csr-1 integrations |
| NCU04869_qPCR_F | TGGTGCGTTCAACCTGAAAA | Used for ChIP-qPCR |
| NCU04869_qPCR_R | TCACCATTTTCTTCCGCACAA | Used for ChIP-qPCR |
| RYq7-ncu06889-F | CATCTTGTATCGAACCCCTTGC | Used for ChIP-qPCR |
| RYq8-ncu06889-R | AAGAGATCGAGGAATGTACGGC | Used for ChIP-qPCR |
| gh3-6-R-qPCR | CAAGAGGCAGGGTGGTGTAAT | Used for ChIP-qPCR |
| gh3-6-F-qPCR | TGCCCATCAATCCCAAGACC | Used for ChIP-qPCR |
| 5 prime landing pad FP | CAACAAGCACATCCAGACACTGAG | Used to create ncu06889::bar |
| 3 prime landing pad RP | CTCTTCTCATGGCCATGTTGATTC | Used to create ncu06889::bar |
| RY49-rtt109-R | CACTTTGTACACACTACACA | Used to genotype rtt109 |
| RY50-rtt109-F | TGTATGGTGAAGCAAAAGCT | Used to genotype rtt109 |
| SUZ12rev | GGGTTGTGCGAGGTGTGG | Used to make tetR-suz12-gfp |
| SUZ12fwd | TCCGTCTTTTGGGTAGCTAGAT | Used to make tetR-suz12-gfp |
| csr1 3p RP w overhang NEW | AATTAATACGACTCACTATAGGGAGGCCCCTGGTTTACTGAGGGC | Used to integrate tetR-suz12-gfp into csr-1 |
| csr1 5 FP overhang | CACTATAGAACTCGAGCAGCTGAAGCAACACCTCCGTCGCCATAAACTCC | Used to integrate tetR-suz12-gfp into csr-1 |
| RY137 Gibson-csr-1-rtt109-3'flank | GATAAGCTTGATATCGAATTCTTAGCATCCTCGTCGACGACCCCTGCCCC | Used to genotype rtt109 inegrations in csr-1 locus |
| RY138 Gibson-csr-1rtt109-5'flank | CACTAGTTCTAGAGCGGCCGCTAAGCTGACGTTTTGGGCTTCTTCCTGAT | Used to genotype rtt109 inegrations in csr-1 locus |
| RTT109 3'UTR | TGTGGGAGAAGGTGATGTTACAAATC | Used to create knockin rtt109-3xflag |
| RTT109-3'CDS-3xflag | CCTCCGCCTCCGCCTCCGCCGCCTCCGCCAGCTGACGTTTTGGGCTTCTTCC | Used to create knockin rtt109-3xflag |
| RTT109-5' CDS-F | AGTCAAGAAACACCAACGGAATTACC | Used to create knockin rtt109-3xflag |
| RTT109-5'UTR-3xflag | TATTCTATAGTGTCACCTAAATAGCTTGGGTCCTCATGATTACTCTTTTCTTTCTTAGTC | Used to create knockin rtt109-3xflag |

| Supplemental Table 2. List of strains used in this study | |  |
| --- | --- | --- |
| Strain # | Genotype | Source |
| S2 | Wild type *N. crassa* (FGSC2489) | FGSC, Kansas City |
| S542 | his-3+::tetO; mus-52::bar+ | 1 |
| S543 | his-3+::tetO; csr-1::p[TrpC]::TetR::eGFP ; mus-52::bar+ | 1 |
| S872 | his-3::tetO, csr-1::p[Dim5]-tetR-suz12-gfp; mus-52::bar+ | This study |
| S866 | ncu06889::bar | This study |
| S858 | rtt109::rtt109-3xflag-hph | This study |
| S8 | mus-52::bar+ | 2 |
| S11 | mus-52::hph+ | 2 |
| S337 | rtt109::hph, mus-51::hph | This study |
| S534 | rtt109::hph | This study |
| S857 | rtt109::hph (Parental) | This study |
| S603 | eed::hph | This study |
| FGSC12340 | rtt109::hph | FGSC, Kansas City |
| FGSC11778 | naf-2::hph | FGSC, Kansas City |
| S539 | asf-1::hph-KD | This study |
| S955 | H3K56R | 3 |
| S957 | H3K56Q | 3 |
| S238 | set-7::hph | This study |
| S960 | eed::hph, ncu06889::bar | This study |
| S961 | rtt109::hph, ncu06889::bar | This study |
| S962 | rtt109::hph Passage 0 | This study |
| S963 | rtt109::hph Passage 1 | This study |
| S964 | rtt109::hph Passage 2 | This study |
| S965 | H3K56R Passage 0 | This study |
| S966 | H3K56R Passage 3 | This study |
| S967 | rtt109::hph, mus-51::hph, csr-1::rtt109-3xflag | This study |
| S968 | rtt109::hph, mus-51::hph, csr-1::rtt109-D145A-3xflag | This study |
| S969 | rtt109::hph, mus-51::hph, csr-1::rtt109-D145A-DD304-305AA-3xflag | This study |
| S970 | rtt109::hph, his-3::tetO, csr-1::p[Dim5]-tetR-suz12-gfp | This study |

1- Tinh-Suong Nguyen, Eugene Gladyshev. Fungal Genetics and Biology. 2020/03/01. DOI: 10.1016/j.fgb.2019.103316.

2-Yuuko Ninomiya. et al. Proceedings of the National Academy of Sciences. 2004-8-17. DOI: 10.1073/pnas.0402780101.

3-Zhang, Chengcheng. et al. Nucleic Acids Research. 2022/04/22. DOI: 10.1093/nar/gkac196.

| Supplemental Table 3. Antibodies used in this study | | | |
| --- | --- | --- | --- |
| Antibody | Host | Supplier | Catalog number |
| H3K27me3 | Mouse | Diagenode (Novus) | NBP2-59295 |
| Anti-FLAG M2 | Mouse | Sigma | F1804-200UG |
| H3K27me3 | Mouse | Abcam | ab6002 |
| H3K36me3 | Rabbit | Abcam | ab9050 |
| Histone H3 | Rabbit | Active Motif | 61799 |
| Anti-mouse IgG (H+L) HRP Conjugate | Goat | ThermoFisher | 31450 |
| Anti-Rabbit IgG HRP Conjugate | Goat | Active Motif | 15015 |
| H3K27me3 | Rabbit | Cell Signaling Technology | C36B11 |
| H3K9me3 | Rabbit | Active Motif | 39161 |
